# Synthetic circRNAs employ IRES activity for translation in cells and in cell-free translation systems

**DOI:** 10.64898/2026.03.28.715045

**Authors:** Philipp Koch, Frederic S. W. Arendrup, Chaehee Lim, Samyukta Narayanan, Amina Noraddin, Massimiliano Clamer, Anders H. Lund, Chun-Kan Chen, Kathrin Leppek

## Abstract

Gene regulation through translation is critical for spatiotemporal protein expression. Internal ribosomal entry sites (IRESes) mediate mRNA-specific translation by recruiting ribosomes to 5′ untranslated regions. Circular RNAs (circRNAs), naturally occurring and stable RNAs, are increasingly used as synthetic tools for sustained therapeutic protein translation by IRES-driven initiation. However, the functionality of different IRESes in synthetic circRNAs remains sparsely characterized. We systematically examine circRNA reporter translation by viral and cellular IRESes in human cells and in diverse *in vitro* translation systems. Improved circRNA purification by urea-PAGE and RNase R-treatment removes contaminants that induce RNA sensing. Viral CVB3 and HCV, as well as cellular *Hoxa9, Chrdl1, Cofilin* and *c-Myc* IRESes, effectively drive circRNA translation. We also establish circRNA translation in a recently developed human cell-free extract that recapitulates IRES-dependent regulation, and allows for precise engineering of HCV IRES-mediated translation. These findings inform IRES selection for synthetic circRNA translation relevant for circRNA-based medicine.

**GRAPHICAL ABSTRACT:** 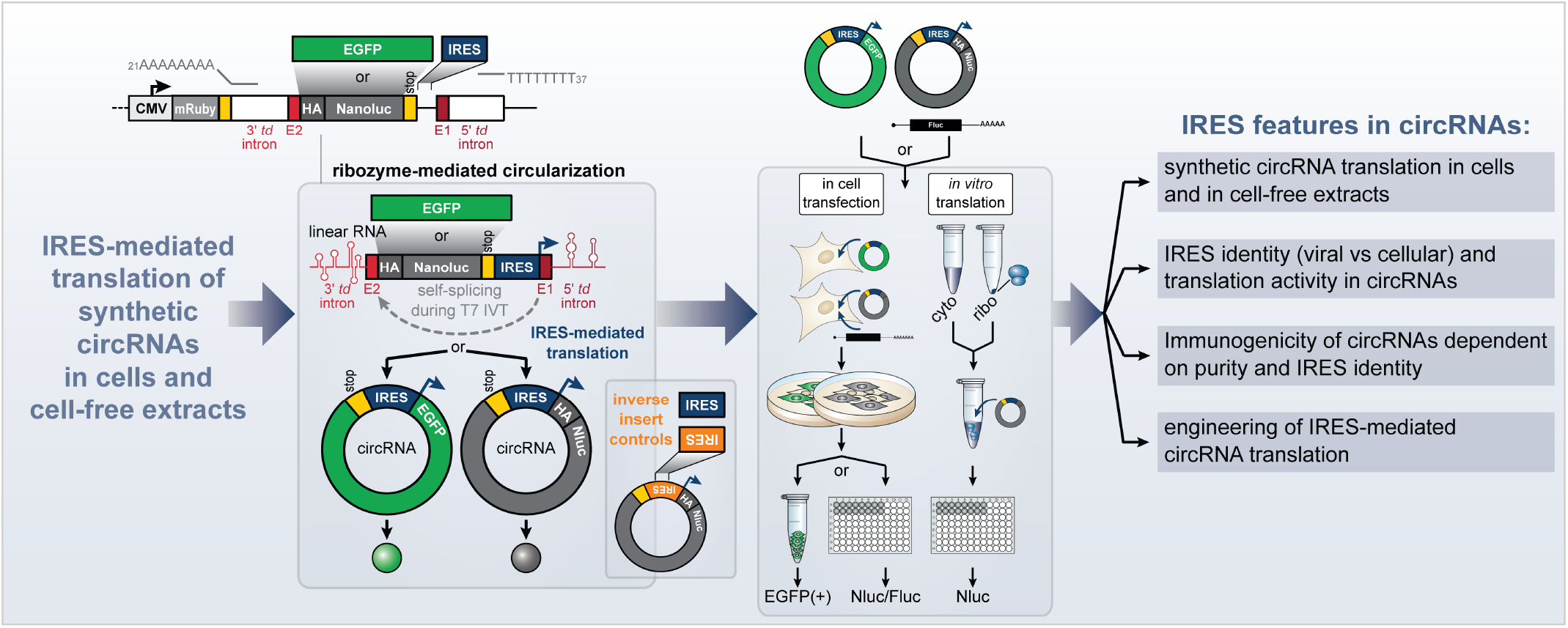

## INTRODUCTION

Recently emerging RNA therapeutics are rapidly becoming a new field of customizable modern medicine (reviewed in^1–5^). The interest to design and optimize mRNA therapeutics is based on the worldwide success of the COVID-19 mRNA vaccines^6,7^, and the huge potential for targeted cancer therapies^8^. Its revolutionary pace in part stems from the fact that RNA medicines can be systematically designed and synthesized more readily at scale compared to other biologics and platforms exist to optimize individual features of mRNA medicine^9–16^. The next exciting wave after vaccines is the circularization of protein-coding mRNAs into single-stranded circular RNAs (circRNAs) as longer-lived versions of linear mRNAs^17–21^. The ends of an mRNA are its crucial destabilizing features attacked by exonucleases. Thus, “endless” circRNAs with covalently linked ends, generated by cellular back-splicing or by *in vitro* self-splicing, are long-lived RNAs^22^. CircRNAs have already been used as an mRNA vaccine against SARS-CoV-2 in rhesus macaques, for example^23^. Rapid antigen translation form and decay of linear mRNA is ideal for vaccination. In turn, circRNAs provide sustained translation useful for protein replacement therapy^24^. circRNAs are already exploited for sustained translation of therapeutic biologics^25,26^. Recent optimization steps^11,12,27,28^ include improved synthesis and circularization of long ORF-encoding mRNAs into circRNAs^29,30^, increased protein yields by several hundred-fold from engineered circRNAs^11^, and chemotopological engineering of circRNAs to attach a 7-methylguanosine (m^7^G)-cap structure to circRNAs for efficient translation^31^. However, these approaches result in protein overexpression. We set out to develop synthetic circRNA systems that allow the study of physiological scales of translation activity. Such activities can range from very high in case of viral internal ribosomal entry sites (IRESes) due to infection requiring high levels of protein synthesis, to lower, context-dependent activity in case of highly regulated elements in the untranslated region (UTR) such as IRESes from cellular mRNAs. Reduced expression is relevant for protein replacement therapy and to conserve the context-dependence of an IRES instead of potential harmful protein overproduction of constitutively active viral IRESes. We aim to study IRES function in circRNAs with the goal to study their mechanism of translation.

Due to the lack of 5′ and 3′ ends in circRNAs, there is no m^7^G at the 5′ cap as in linear mRNAs required for cap-dependent initiation. On mRNAs, the eukaryotic initiation factor F (eIF4F) complex at the m^7^G-cap recruits the pre-initiation complex for 5′ UTR scanning and start site recognition^32^. circRNAs initiate translation fundamentally differently and rely on cap-independent mechanisms to recruit ribosomes. Currently, internal translation initiation on circRNAs is thought to occur via two alternate mechanisms: 1. The incorporation of the RNA modification N^6^-methyladenosine (m^6^A) in the UTR, which binds to the eIF3 complex to recruit the 40S ribosome^11,12,33^, and 2. Internal RNA structural ensembles that recruit initiating ribosomes (IRESes). mRNAs can harbor diverse functional regions in UTRs that regulate their translation and stability^34,35^. IRESes are one such category of often structured, functional 5′ UTR elements that are found in many viral^36,37^ and certain eukaryotic mRNAs^12,38,39^. They facilitate alternate, cap-independent translation, bypassing most required steps of conventional initiation^32,37,40–42^. Ribosome-contact to IRESes can occur directly or through RNA-binding proteins (RBPs), so called IRES *trans*-acting factors (ITAFs)^43^, and via direct base pairing or tertiary interactions with ribosomal RNA (rRNA)^44–46^. “Cellular IRESes” are IRESes that occur in 5′ UTR of endogenous transcripts that can act as translational enhancers that bind ribosomes independently from the cap. Initial examples were often discovered because they enabled continued translation of select mRNAs upon cellular stimulation or stress, conditions that repress global cap-initiation and IRES-mediated translation was sustained (reviewed in^40,43,47–49^). Systematic screens^12,38,39^ showed that cap-independent activity is estimated to exist in ∼10% of human 5′ UTRs^38^ and is dynamic across cell states^38,50^. Mammalian IRESes can contribute to translation even during ongoing cap-mediated translation^51^. Examples include a subset of *Homeobox* (*Hox*) mRNAs that contain structured IRES-like elements^52^, of which *Hoxa9* uses a short stem-loop to directly recruit the 40S ribosome via the rRNA expansion segment ES9S in 18S rRNA^44^. These IRES-like translation enhancers are active under physiological conditions in development in concert with a potent cap-repressor in the same 5′ UTR^52^, a means to more specifically control spatiotemporal translation. One of the first cellular IRES examples was found in the *Drosophila Antennapedia* 5′ UTR^53,54^, a fly Hox gene homologous to the vertebrate HOXA cluster. circRNAs only have one UTR and IRES activity of this UTR serves as the ribosomal entry point^12,30,55^. Endogenous IRES-containing circRNAs that encode a protein were first described 30 years ago^56^. Several examples of naturally occurring back-spliced circRNAs with physiological functions in human health and disease have been described^19,57–64^. But only a few circRNA-encoded proteins are known in mammals^28,65–68^, and their potential IRESes remain mostly unexplored.

For therapeutic applications of synthetic circRNAs, the immunogenicity of RNA^25,29,69–72^ is a key feature. However, it is unknown whether the IRES identity affects the degree of immune sensing of circRNAs. Retinoic acid-inducible gene-I (RIG-I) and melanoma differentiation-associated gene 5 (MDA5) are key cytoplasmic innate immune sensors of foreign RNAs. RIG-I is activated by short 5′-triphosphate RNA structures^73^, and MDA5 recognizes long double-stranded (ds)RNAs^73,74^. Neither feature is typically present in circRNAs lacking a 5′ end. Previous work showed that ribozyme intron origin determines the level of self from non-self circRNA immune recognition, with exogenous circRNAs being sensed by RIG-I^29,72,75^. RNA circularization diminishes immunogenicity caused by triphosphorylated, linear RNA contaminants in the RNA preparation^25^. Cell-foreign intron-derived circRNAs used in this study can induce a stronger RIG-I immune response^29,72^ than self intron-derived circRNAs that confer self-identity^76^, possibly due to *td* intron scar sequences without associated host splicing factors, and m^6^A installation during splicing by endogenous introns as a “self” marker^69^. RIG-I was found to sense foreign *td* intron-derived circRNA in HEK293T cells^29^. Sequence fragments in group-I ribozyme intron-derived circRNAs may also form immunogenic dsRNA structures or distort circRNA folding that inhibit activation of the dsRNA receptor protein kinase R (PKR)^70,72^. Thus, both were reported: exogenous circRNAs provoking a strong RIG-I-mediated innate immune response^29^, and no activation of several RNA sensors^25^. Different findings on circRNA immunogenicity likely reflect differences in circRNA preparation and assay conditions^27,71^. In addition, it is reasonable that dsRNA structures within IRESes, beyond sequence motifs, need to fold into stable structures inside cells to recruit ribosomes, which in turn may bind and activate dsRNA sensors. Thus, we focus on the immune sensing across diverse IRESes in circRNAs.

Understanding circRNA translation and identifying optimal IRESes for sustained protein expression remains a major challenge. The mechanisms of IRES-driven circRNA translation, the role of IRES origin, and their immunogenicity are not well understood. We tested viral and cellular IRESes in synthetic circRNA reporters produced by ribozyme activity, examining them in both human cells and cell-free systems. Using urea-PAGE gel extraction, we assessed how circRNA purity influences the ability of different IRESes to affect RNA sensor activation. We demonstrated that a HEK293T cell-free system more accurately mimics physiological IRES-dependent translation than commonly used extracts, and allows circRNA translation to be engineered through targeted mutations in the ribosome-interacting regions of the HCV IRES. Overall, synthetic circRNA translation may be constrained by ITAF dependence and the preservation of their regulation across in-cell and *in vitro* systems, as well as the RNA processing history of circRNAs. Together, by combining reporter and IRES features for immune-silent and enhanced circRNA translation in cells and across *in vitro* extracts, we enable the customization of circRNA-derived protein yields depending on the selection of specific IRESes, allowing circRNAs to be used to investigate IRES activity and translation mechanisms in diverse cellular and *in vitro* settings.

## MATERIAL & METHODS

### Cell Culture and Transfection

Human HEK393T (ATCC: CRL-3216) were cultured in Dulbecco’s Modified Eagle’s Medium (DMEM, Gibco, 11965-118) containing 2 mM L-glutamine, supplemented with 10% fetal calf serum (Gibco, ES009-B), 100 U/ml penicillin and 0.1 mg/mL streptomycin (EMD Millipore, TMS-AB2-C or Gibco, 15140-122) (regular medium, RM) at 37°C in 5% CO_2_-buffered incubators.

For luciferase assays following RNA transfection, ∼1.5 X 10^5^ HEK393T cells were seeded per well in 24-well dishes and transfected the following day with 1 μg of circRNA, generated by IVT (NEB HiScribe T7 kit, E2040S) and subsequently purified using RNase R (Biozym, 172010) digestion with or without prior 4% urea-PAGE purification, and 100 ng linear capped and polyadenylated control *hHBB*-Fluc mRNA (Firefly Luciferase reporter) using 5 μL Lipofectamine Messenger Max (Invitrogen, LMRNA001) and Opti-MEM (Gibco, 11058-021) according to the manufacturer’s instructions in serum-free and antibiotic-free DMEM. The medium was changed to regular DMEM 6-7 hrs after transfection and cells were collected 24 hrs post-transfection (hpt). The IVT and capping and polyadenylation procedure for the reference linear *hHBB*-Fluc mRNA and *hHBB*-Nluc mRNA have been described elsewhere^10^. Both reference linear mRNAs have both hHBB 5′ and 3′ UTRs and were generated using the respective mRNA-encoding pGL3 plasmid templates.

For luciferase assays following Tornado plasmid transfection, ∼0.8 X 10^5^ HEK393T cells were seeded per well in 24-well dishes and transfected the following day with total 1 μg plasmid DNA using 1 μL Lipofectamine 3000 and 1 μL P3000 (Invitrogen, L3000001) and Opti-MEM (Gibco, 11058-021) according to the manufacturer’s instructions in serum-free and antibiotic-free DMEM. The medium was changed to regular DMEM 12 hrs after transfection and cells were collected 4 days post-transfection. An additional media change was performed 3 dpt.

For immune sensing experiments, ∼1.5 X 10^5^ HEK393T cells were seeded per well in 24-well dishes and transfected the following day with 100-500 ng of circRNA (generated by IVT (NEB HiScribe T7 kit, E2040S) and subsequently purified using RNase R (Biozym, 172010) digestion with or without prior 4% urea-PAGE purification) using 2.5 μL Lipofectamine Messenger Max (Invitrogen, LMRNA001) and Opti-MEM (Gibco, 11058-021) according to the manufacturer’s instructions in serum-free and antibiotic-free DMEM. The medium was changed to regular DMEM 6-7 hrs after transfection and cells were collected 24 hrs post-transfection.

For EGFP-based FACS readouts, ∼1.5 X 10^5^ HEK393T cells were seeded per well in 24-well dishes and transfected the following day with 500 ng - 2 μg of circRNA (generated by IVT (NEB HiScribe T7 kit, E2040S) and subsequently purified using RNase R (Biozym, 172010) digestion using 2.5 μL Lipofectamine Messenger Max (Invitrogen, LMRNA001) and Opti-MEM (Gibco, 11058-021) according to the manufacturer’s instructions in serum-free and antibiotic-free DMEM. The medium was changed to regular DMEM 6-7 hrs after transfection and cells were collected 24 hrs post-transfection.

### Plasmid Constructions

The following plasmids have been described previously: mRuby3-ZK-spEGFP^12^ was kindly provided by Chun-Kan Chen (Washington University School of Medicine, St. Louis); IRES-containing derivatives of mRuby3-ZK-spEGFP^81^; and pcDNA3.1-5′UTR-3xHA-NLuc (pKL401)^110^.

For cloning of the Tornado translation system plasmids, the pcDNA3.1+-Tornado-split-nLuc plasmid backbone (addgene ID 212611) was used for all subsequent cloning. The tested sequences were PCR-amplified from plasmids with accordingly designed Gibson assembly overhangs using primers PK_352–PK_391 according to manufacturer’s recommendations (NEB Q5 Polymerase, M0493S). The plasmid backbone was digested using *EcoRI*-HF (NEB, R3101S) and *BsiWI*-HF (NEB, R3553S) restriction enzymes according to manufacturer’s instructions. The Gibson Assembly reaction was performed using the NEBuilder HiFi DNA Assembly Master Mix (NEB, E2621L) with recommended insert:backbone ratios according to the varying insert lengths. This resulted in plasmid series pPK055-pPK074.

For cloning of the plasmid for circRNA *in vitro* transcription, the mRuby-circEGFP-ScaI-v3 plasmid (pKL480), kindly provided by Chun-Kan Chen, was digested using the EcoRV-HF (NEB, R3195S) and ScaI-HF (NEB, R3122S) restriction enzymes according to manufactures instructions were used for cloning constructs pPK001-005, pPK042, pPK043, pPK044-074, pPK048, pPK076-079. The excised EGFP-encoding fragment was replaced by an 3xHA-NanoLuciferase reporter (3xHA-Nluc) encoding reporter using the NEBuilder HiFi DNA Assembly Master Mix (NEB, E2621L) with recommended insert:backbone ratios, leading to the pPK001 empty vector. The pcDNA3.1-5′UTR-3xHA-NLuc plasmid was used as the PCR template for amplification of the 3xHA-Nluc reporter. Next, the tested IRES sequences were cloned into the mRuby-circ3xHA-NanoLuc plasmid (pPK001) after plasmid linearization by *ScaI* digestion using the Gibson assembly (NEB, E2621S). the tested IRES sequences were PCR amplified from plasmids using primers PK_001-PK_010, PK_268-PK_281, PK_438-441 according to manufacturer’s recommendations (NEB Q5 Polymerase, M0493S). After digestion, the digested backbone plasmid and the PCR-amplified IRES inserts using the indicated primers were purified by agarose gel extraction according to manufacturer’s instructions (NEB, T1020S). Plasmids were used for subsequent *in vitro* circRNA generation.

After transformation of NEB 5-alpha competent *E. coli* (High Efficiency) bacteria (NEB, C2987I) and following Midiprep (QIAGEN Plasmid Plus Midi Kit, 12945), all plasmid sequences were verified by DNA sequencing (Eurofins Genomics or Microsynth). All oligonucleotides were purchased from IDT. A list of all plasmids (**Appendix Table S1**) and primers (**Appendix Table S2**) are listed in the Appendix.

### Cell Harvesting, Luciferase Activity Assay, and RNA Isolation

Cells were incubated for 2 min at 37°C with 50 μL 0.05% Trypsin-EDTA (Thermo Fisher Scientific B3026-15400054). Reaction was stopped by adding 450 μL regular medium and cells were transferred into 1.5 mL Eppendorf tubes (Eppendorf 211-2164). After centrifugation at 600 g for 5 min at RT, cells were washed with 500 μL DPBS (Thermo Fisher Scientific B3026-14190169). The DPBS was removed and the cell pellet was used for subsequent analysis.

For luciferase measurements, cells were resuspended in 50 μL 1x Passive Lysis Buffer (Promega, E1941) and incubated for 30 min at 1000 rpm at RT. Debris was removed by centrifugation at 10.000 g for 5 min at RT and 40 μL of cell lysate was transferred into a Nunc MicroWell 96 well plate (Thermo Fisher Scientific, 137101). Fluc and Nluc expression was quantified using the Nano-Glo Dual Luciferase Reporter Assay (Promega, N1610) and the Promega GloMax Explorer. Luciferase reporter activity is expressed as a ratio between Nluc and Fluc (Nluc/Fluc).

For FACS measurement, cell pellets were resuspended in a volume of 180 μL FACS buffer (1xPBS, 1% BSA, 0.1% NaN_3_). The EGFP signal intensity was then measured by using the ABI Attune Acoustic Focusing Cytometer (Thermo Scientific). FACS data were exported as FCS files and further analyzed by FlowJo v10.8 (BD). Viable cells and singlets were identified based on the FSC and SSC channels. Finally, the EGFP signal was measured and used for calculation of the median fluorescence intensities (MFIs).

For analysis of circRNA immunogenicity, cell pellets were resuspended in 500 μL TRIzol (Thermo Fisher Scientific, 15596026). Total RNA isolation was performed by adding 100 μL Chloroform, vortexing and centrifugation at 12.000 g for 25 min at 4°C. The aqueous phase was collected and RNA was extracted using the Zymo Clean & Concentrator-5 column (Zymo Research, R1016) according to the manufacturer’s protocol. The RNA was eluted in 34 μL RNase-free water and DNA was removed by Turbo DNaseI treatment (Thermo Fisher Scientific, AM2239) (2 μL/sample) and incubated for 30 min at 37°C. After DNA degradation, the RNA isolation procedure using the Zymo Spin Column I was repeated and final RNA concentration was quantified by Nanodrop measurement.

### Northern Blot Analysis of EGFP-encoding circRNAs

Northern Blot analysis of EGFP encoding circRNAs was performed as described in^12^. In brief, four 15 cm plates of HEK293T cells were transfected with diluted mRuby-ZKSCAN1-split-EGFP reporter plasmids using Lipofectamine 3000 Transfection Reagent (Thermo Fisher Scientific). Transfected cells were grown for five days, trypsinized, and sorted by EGFP signal intensity. mRuby+/EGFP+ cells were collected and total RNA was purified using the Quick-RNA Midiprep Kit (Zymo Research). 50 mg of total RNA was treated with 500 U RNase R (MACLAB:RNASR-200) at 37°C for 30 min in 1X RNase R Buffer (0.2 M Tris-HCl (pH 8.0), 1 mM MgCl_2_ and 1 M LiCl). Another 50 mg of total RNA was incubated in 1X RNase R Buffer without RNase R addition at 37°C for 30 min (RNase R(-) sample). Total RNA with or without RNase R treatment was purified using RNA Clean & Concentrator-25 (Zymo Research: R1017), respectively. Northern blotting was then performed using theNorthern Max-Gly Kit (Thermo Fisher Scientific: AM1946) according to the manufacturer’s protocol. Specifically, purified RNA was incubated with equal volume of Glyoxal Load Dye (Thermo Fisher Scientific: AM8551) at 50°C for 30 min, loaded into 1% agarose gels made in 1X Gel Prep/Gel Running buffer (Thermo Fisher Scientific: AM8678), and ran at 75 V for 60 min. Gels were then stained in 1X SYBR Gold Nucleic Acid Gel Stain (Thermo Fisher Scientific: S11494) diluted in 1X Gel Prep/Gel Running buffer at room temperature for 5 min, and transferred to BrightStar-Plus Positively Charged Nylon Membranes (Thermo Fisher Scientific: AM10100) using the Trans-Blot Turbo Transfer System (BioRad) with STANDARD SD program for 30 min. We then crosslinked the blots with a Stratalinker 1800 (Stratagene) using autocrosslink mode. RNA ladders on the blots (Millennium RNA Markers, Invitrogen: AM7150) were visualized and marked on a E-Gel Imager (Thermo FisherScientific). The blots were blocked in ULTRA hyb Buffer (Thermo FisherScientific: AM8670) pre-warmed to 68°C with 20 U SUPERaseIn RNase Inhibitor (Invitrogen: AM2694) at 65°C for 30 min. 0.1 nM biotinylated ssDNA probes synthesized by IDT against the corresponding region on the reporter RNA (EGFP junctional probe: /5BiosG/GTAGTGGTCGGCGAGCTGCACGCTGCCGTCCTCGATGTTGTGGCGGATCTTGAA GTTCAC) were added to the blots, respectively, in ULTRAhyb Buffer and incubated at 65°C overnight. The blots were washed with Northern Blot Wash Solutions (Thermo Fisher Scientific: AM8673), followed by blocking in EveryBlot Blocking Buffer (BioRad: 12010020) at room temperature for 40 min. After the blocking, the blots were incubated with IRDye 800CW Streptavidin (LI-COR Biosciences; 1:200) in PBS at room temperature for 40 min, washed with Northern PBS. The blots were imaged with Odyssey M Imaging System (LI-COR Biosciences).

### cDNA Synthesis and Quantitative RT-PCR (RT-qPCR) Analysis

Cells transfected with 100–500 ng circRNA were collected in 500 μL TRIzol (Invitrogen, 15596). Total RNA was isolated from the aqueous phase using Zymo Clean & Concentrator-5 column (Zymo Research, R1016) and treated with TURBO DNaseI (Ambion, AM2238) followed by a second Zymo column purification step. For reverse transcription-quantitative PCR (RT-qPCR) analysis, cDNA was synthesized from 300-500ng of total RNA using iScript Supermix (Bio-Rad, 1708840) containing random hexamer primers, according to the manufacturer’s instructions. qPCR reactions were assembled in 384-well plates using 2.5 μL of a 1:4-1:5 dilution of a cDNA reaction, 300 nM of target-specific primer mix and the Powertrack SYBR MM (Thermo Fisher Scientific, A46111) or 5x EvaGreen qPCR-Mix (Bio-Budget, 80-5820000) in a final volume of 10 μL per well. SYBR green detection-based qPCR was performed on a CFX384 machine (Bio-Rad) or QuantStudio qPCR machine (Thermo, applied biosystems, 15721248). Data was analyzed and converted to relative RNA quantity manually or using CFX manager (Bio-Rad). Human GAPDH mRNA was used as the internal normalizer in all RT-qPCR analyses.

### *In vitro* Transcription of circRNAs and RNase R Purification

For circRNA IVT, plasmids pPK001-pPK005, pPK042, pPK043, pPK044–074, pPK048, pPK076–079 were used as PCR templates using the AccuPrime Pfx DNA Polymerase (Thermo, Invitrogen, 1876525) in order to prepare the T7 promoter containing *in vitro* transcription templates (5 μL 10x AccuPrime Pfx Reaction Mix, 1.5 μL forward and reverse primer 10 μM (LL078, LL079), 0.5 μL Pfx Polymerase, 1.5 μL DMSO, 0.5 μL MgSO_4_ (50 mM) in a total reaction volume of 50 μL (protocol according to manufactures guidelines). The PCR products were purified using the Monarch PCR & DNA Cleanup kit (NEB, T1030L) and 1 μg DNA was subsequently used for *in vitro* transcription using the High Scribe T7 High Yield RNA synthesis Kit (NEB, E2040S) according to manufacturer’s protocol (20 μL total reaction volume) supplemented with 40 U RiboLock RNase inhibitor (Thermo, EO0381) and incubated for 2 hrs at 37°C and 800 rpm. The 20 μL sample was diluted with 70 μL ddH_2_O and remaining DNA was digested using 2 μL TurboDNaseI (Life Technologies, AM2238, 20 min incubation at 37°C at 500 rpm). The enriched RNA was purified using the PureLink RNA Mini Kit (Thermo, 12183018A) and non-circularized RNA was subsequently removed by RNase R (Biozym, 172010) digestion (working concentration 1 U/μg RNA, incubation at 37°C for 30 min at 500 rpm) for the non-gel purified circRNA samples. For gel purified and RNase R-digested circRNAs, unpurified IVT RNA was supplemented with 2x RNA loading dye (Thermo Fisher Scientific, R0641), loaded onto a 4% urea-PAGE gel (up to 7 μg total RNA was loaded per lane in order to achieve optimal separation between the circRNA and the non-circular contaminating RNA species) and the gel was run for up to 3.5 hrs at 180 V (dependent on the circRNA length). Total RNA was visualized using SYBR Gold (Thermo Fisher Scientific, S11494). The circRNA band was identified by comparing the sample lanes with the control lane containing a small amount of already digested circRNA of the same RNA species. The circRNA was cut out and stored in crush soak buffer (200 mM sodium chloride, 10 mM Tris-HCl pH 7.5, 1 mM EDTA pH 8.0) at -80°C overnight. On the next day, gel pieces were thawed and incubated for 2.5 hrs at RT under constant rotation. Next, 300 μL crush soak buffer was supplemented with 700 μL ethanol and RNA precipitation was performed at -20°C overnight. RNA was finally pelleted at 4°C 18.000 g for 30 min, ethanol was removed and the pellet was air dried for 15 min at RT. RNA was resuspended in 25 μL RNase-free water and potential non-circularized RNA was subsequently removed by RNase R (Biozym, 172010) digestion (working concentration 0.3 U/μg RNA, incubation for 7 min at 37°C at 500 rpm). The PureLink RNA Mini Kit was used for circRNA purification after RNase R treatment. In order to verify the successful linear RNA degradation by RNase R treatment, 10 μg of IVT RNA was treated as described beforehand with the exception of the RNase R addition. 500 ng of this negative control were loaded together with 500 ng of the digested sample on a formaldehyde gel to verify the complete degradation of the linear RNA. Alternatively, the purity of the circRNA was assessed for more accurate quantification using the Agilent Technologies 4200 TapeStation system with the High Sensitive RNA Screen Analysis test (Agilent Technologies, Cat. No. 5067-5579).

### *In vitro* Translation Systems (Wheat Germ Lysate (WGL), Rabbit Reticulocyte Lysate (RRL), Tripartite cell-free translation system & Immagina ReCet HEK293T system)

*In vitro* generated and purified circRNA was used throughout all *in vitro* translation systems. For the Promega WGL (L4380) and RRL (L4960) extracts, 10-500ng of linear Fluc mRNA and Nluc-encoding circRNA were used at a ratio of 1:10. Using Promega’s RRL, the provided lysates were thawed on ice according to manufacturer’s instructions, reactions were assembled (total volume 25 μL, including 17.5 μL RRL) and incubated for 90 min at 30°C at 300 rpm. Next, the reactions were diluted using 25 μL 1x Passive Lysis Buffer (Promega, E1941) and Nluc/Fluc expression was detected as described before. Using Promega’s WGL, the provided lysates were thawed on ice according to manufacturer’s instructions, reactions were assembled (total volume 25 μL, including 12.5 μL WGL, 1 μL KAc, 0.5 μL RNasin (Promega, B2775-N2615)) and incubated for 2 hrs at 25°C at 300 rpm. Nluc/Fluc expression was detected as described before using the Nano-Glo Dual Luciferase Reporter Assay (Promega, N1610) and the Promega GloMax Explorer. The tripartite cell-free translation system was prepared according to Arendrup *et al*., (2025)^82^ using HEK293T cells with the following specifications: Cytoplasmic lysates were generated by lysing the cells in hypotonic lysis buffer and and multiple rounds of application of shearing forces using a prechilled 27G needle. After each pipetting step, the integrity of the cells was checked under the microscope in order to avoid over-lysing the cytoplasmic preparation. The subsequent steps were performed following the recommendations of Arendrup *et al*. (2025)^82^: The ribosomal fraction was also prepared according to the described protocol using a short puromycin pre-treatment before lysing the HEK293T cells. An additional MNase treatment after isolation was not performed in order to provide maximum translation activity of the prepared ribosomes, keeping in mind that this decision could lead to off-target translation due to endogenous mRNA contamination. Final reactions (10 μL total) were pipetted on ice and included 3 μL translation buffer, 5 μL cytoplasmic lysate, 1 μL ribosomes and 1 μL RNA (90 ng). The ribosomal fraction was added last and was pipetted in parallel into the samples in order to ensure the same incubation time across samples. The reaction was incubated for 30 min at 37°C and stopped by adding 40 μL ice cold DPBS. Nluc/Fluc expression was detected as described before using the Nano-Glo Dual Luciferase Reporter Assay (Promega, N1610). The Immagina ReCet HEK293T *in vitro* translation system was used according to manufactures instructions. Provided chemicals and lysates were thawed on ice for at least 40 min and a master mix (MM) containing 10x Translation Buffer, RNase Inhibitor, Saline solution 1 & 2, Cytoplasmic extract, ATP and Ribosome extract was generated. Final reactions were assembled on ice adding the 8.5 μL MM last, in order to start the translation reactions simultaneously. Total reaction volumes were adjusted to 24 μL using 50-400ng purified Nluc encoding circRNA (urea-PAGE gel purified and RNase R and column purified) or 50-400ng linear capped Fluc control RNA. Reactions were incubated for 1 hr at 37°C. Reactions were stopped by adding 26 μL cold DPBS. Nluc/Fluc expression was detected as described before using the Nano-Glo Dual Luciferase Reporter Assay (Promega, N1610) and the Promega GloMax Explorer.

### Western Blot Analysis and Antibodies

Proteins were resolved on self-made 4-20% or 15% polyacrylamide gradient Tris-glycine SDS-PAGE gels and transferred onto 0.2 μm pore size PVDF membranes (Biorad) using the semi-dry Trans-Blot Turbo system (Biorad, 170-4273). Membranes were then blocked in 1x PBS-0.1% Tween-20 containing 5% non-fat milk powder for 1 hr, incubated with primary antibodies diluted in the same solution for 1 hr at RT or overnight at 4°C, and washed four times for 5 min in 1x PBS-0.1% Tween-20, incubated with secondary antibodies for 1 hr in 1x PBS-0.1% Tween-20 and washed four times for 15 min in 1x PBS-0.1% Tween-20. Horseradish peroxidase (HRP)-coupled secondary antibodies (anti-mouse, GE Healthcare) in combination with Clarity Western ECL Substrate (Biorad, 170-5061) and imaging on a ImageQuant 800 (Amersham, Cytiva) were used for detection. Antibodies were diluted in 1x PBS-0.1% Tween-20 at 1:1000-1:3000 dilution in 5% BSA (w/v) in 1x PBS-A (azide). The following primary antibody was used for Western blot analysis: mouse monoclonal anti-HA antibody (Thermo Fisher Scientific, 26183).

### Quantification and Statistical Analysis

In all figures, data was presented as mean, SD or SEM as stated in the figure legends, and *p ≤ 0.05 was considered significant (ns: p > 0.05; *p ≤ 0.05; **p ≤ 0.01; ***p ≤ 0.001; ****p ≤ 0.0001). Blinding and randomization were not used in any of the experiments. The number of independent biological replicates used for the experiments are listed in the figure legends. Tests, two-tailed unpaired Student’s t-test, if not stated otherwise, and specific p-values used are indicated in the figure legends or figures themselves. In all cases, multiple independent experiments were performed on different days to verify the reproducibility of experimental findings.

## RESULTS

### Ribozyme-mediated generation of synthetic IRES-containing circRNAs

Synthetic circRNAs are a powerful tool to study cap-independent translation initiation. We first aimed to establish a controlled *in vitro* system to generate circRNAs containing an IRES in their UTR. Then, we employed these circRNAs to investigate IRES-dependent activity in cells and *in vitro*, their immunogenicity, and engineering of their activity. We first tested the activity of IRESes of different origins in *in vitro* synthesized circRNAs. Synthetic circRNAs are different from plasmid-derived back-splicing systems, where split-reporter constructs rely on RNA processing and spliceosome activity. Ribozyme-based *in vitro* circRNAs avoid cryptic or artificial promoter effects from IRES inserts that are possible in plasmid-based systems. We used a synthetic approach to study diverse IRESes in *in vitro* generated circRNAs with the potential for long-lived RNA therapeutics (**Fig. 1A**). We generated circRNAs encoding the EGFP or Nanoluciferase (Nluc) reporter ORF as cargo upstream of an IRES during *in vitro* transcription (IVT). RNA circularization is mediated by permuted split group I introns from the phage T4 *thymidylate synthase* (*td*) gene. The 5′ and 3′ split introns fold into an active self-splicing ribozyme through distally extended RNA interactions^77,78^, that relies on magnesium and a free guanosine to initiate splicing (**Fig. 1A**, right panel). This ribozyme has been successfully used for circRNA synthesis previously^20,29,30,77,79,80^. We set out to optimize the efficiency of the ribozyme-mediated reaction, which resulted in the critical improvement of using primers that attach A/T-tails to DNA templates to drastically increase the circularization efficiency (**Fig. 1A**). We assumed that the complementary distal ends help circularize the DNA ends and promote intron proximity for enhanced ribozyme cleavage^30^. The resulting circRNA retains a defined E1/E2 exon-exon junction and scar sequence essential for ribozyme activity, between the IRES and the start codon. To ensure consistent processing of the IRES-ORF fusion site, and to avoid effects of IRES identity on ribozyme activity, the IRES was placed downstream of the ORF. This design also prevents that any residual linear RNA after circRNA purification can contribute to reporter expression. Additionally, inverse IRES sequences were included to control for effects of differences in IRES length, structure, and GC-content on ribozyme efficiency. To test diverse IRESes in circRNA reporters, we selected two viral IRESes previously shown to have IRES activity in plasmid-derived back-spliced circRNAs^81^ (**Fig. 1A**, right panel). As test cases, we chose the hepatitis C virus (HCV) and coxsackievirus B3 (CVB3) viral IRESes. The *human hemoglobin subunit beta* (*hHBB*) 5′ UTR, commonly used in RNA therapeutics e.g. COVID19 mRNA vaccines, served as a negative control. The CVB3 IRES has been widely used in circRNAs, including in IVT-derived circRNAs^30^. With the goal to optimize IRES-driven translation in synthetic circRNAs and to investigate IRES features in circRNAs, we applied RNase R, a 3′-to-5′ exonuclease that degrades all linear RNA species with accessible ends, and column-purified synthetic IRES-containing circRNAs to either in-cell transfection or *in vitro* translation assays, summarized in **Fig. 1B**. For all circRNAs, T7 IVT RNA products are treated with RNase R, column-purified to enrich circRNAs, and RNAs are analyzed by gel (denaturing formaldehyde (FA)-agarose) and capillary (TapeStation) electrophoresis (**Fig. 1C, D**; **Fig. S1A, B**). FA-gels and TapeStation analysis showed both the linear RNA and circRNA as IVT products for EGFP (**Fig. 1C**) and Nluc (**Fig. 1D**), with up to ∼50% circRNA for Nluc. Digestion of the IVT RNA with the exonuclease RNase R and column purification removed most linear RNA. circRNAs were enriched as a single band, independent of insert identity, on FA gels (**Fig. 1C; Fig. S1A**) and TapeStation RNA chips (**Fig. 1D; Fig. S1B**). Residual RNase R-resistant *td* introns, stable dsRNA side products of lower molecular weight, and minor linear RNA remained. Remaining linear mRNAs are uncapped and are expected to be unstable and not translated. Thus, reporter translation should mainly reflect internal translation initiation on circRNA. We concluded that synthetic circRNA can be efficiently generated and enriched *in vitro*. We next evaluated specific IRES performance in synthetic circRNAs in cells.

**Figure 1.**
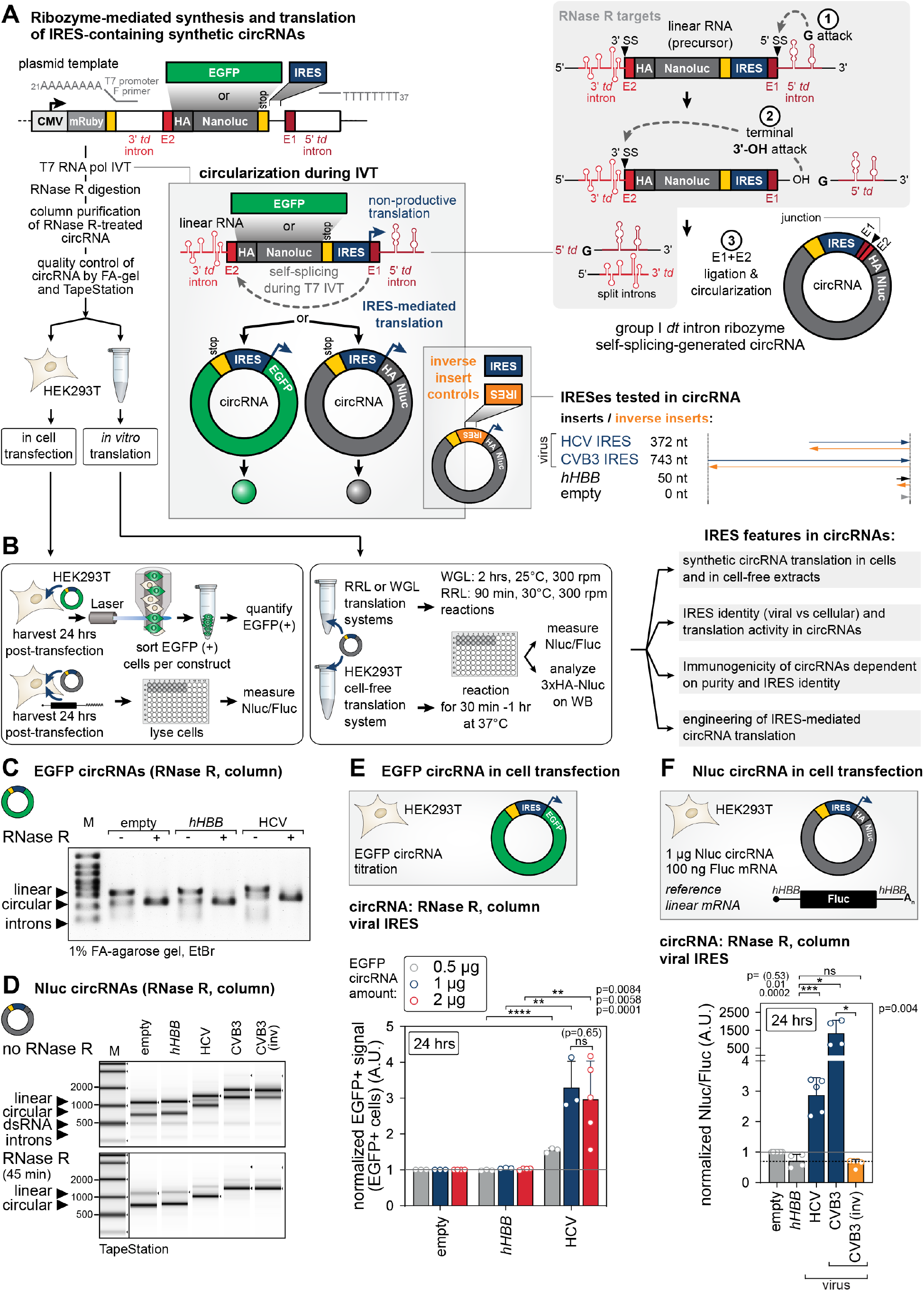
circRNA reporter design, *in vitro* transcription, RNase R purification of synthetic circRNAs, and viral IRES-mediated circRNA-Nluc translation in transfected cells. **(A)** Experimental outline of the plasmid-based *in vitro* generation of synthetic circRNA and reporter assays for evaluation of IRES activity of different IRES inserts, including inverse sequences as controls for insert length and GC-content. Following DNA template synthesis (PCR amplification from plasmid backbone using poly(A)- and poly(T)-including primers), circRNA was generated by T7 *in vitro* transcription. Linear RNA species were removed by RNase R-treatment and column purification. Resulting circRNA was subsequently used for in-cell transfection or for *in vitro* translation systems. EGFP and 3xHA-Nluc reporter systems were used throughout the following analyses. (Right) Illustration of the circularization reaction mediated by group I *dt* introns leading to self-spliced circRNA. The observed sequence scar formed by the remaining parts of the *td* introns are highlighted in light and dark red. Remaining split introns were generated as side products and need to be removed. (Right, bottom) Overview of the tested IRES sequences indicating length and viral origin of the tested IRESes and included controls.**(B)** Experimental outline of the assays using *in vitro* generated circRNA reporters encoding EGFP or Nluc after in-cell transfection into human HEK293T cells (left), or in cell-free *in vitro* translation systems (right). EGFP-encoding circRNAs are indicated as green circles, Nluc-encoding circRNAs are indicated as grey circles. Nluc-encoding circRNA was co-transfected with linear capped *hHBB*-Fluc mRNA used as a transfection control. Cells were harvested 24 hrs post transfection (hpt). Subsequent *in vitro* translation readouts, incubation times and reaction conditions of the different extracts are indicated. We tested four different aspects or features based on IRES identity in circRNAs.**(C)** Quality control of the generated EGFP reporter including circRNAs using 1% FA-agarose gel. Linear side products (upper band) disappear by RNase R digestion. Only the circular RNase R-resistant band remains (lower band). 500 ng total RNA was loaded per lane. The RiboRuler High Range RNA Ladder (Thermo) serves as a reference.**(D)** Quality control, using the High Sensitivity RNA ScreenTape (Agilent), of the generated Nluc circRNAs before purification (upper panel) and after RNase R digestion and column purification (lower panel). Remaining contaminants can be observed as light grey bands.**(E)** Optimal RNA concentration was evaluated using the EGFP-encoding circRNAs 24 hpt. RNA was purified by RNase R digestion and column purification. 0.5, 1, and 2 μg circRNA was transfected and evaluated using EGFP signal intensities normalized to the empty vector control. Calculated median fluorescence intensities (MFIs) of EGFP are shown. Bar graphs are indicating mean values ± SD, n = 3-5. Empty vector control was normalized to 1; ns, not significant.**(F)** IRES activity of the tested sequences was evaluated 24 hpt using Nluc circRNAs. RNA was purified by RNase R digestion and column purification. Tested sequences are depicted in dark blue, inverse controls in orange. 1 μg of circRNA was co-transfected with 100 ng hHBB-Fluc linear capped control RNA. Nluc data were normalized to Fluc transfection control and Nluc/Fluc ratios are normalized to empty control; n= 3-5. In all figures, data was presented as mean ± SD as stated, and *p ≤ 0.05 was considered significant (ns: p > 0.05; *p ≤ 0.05; **p ≤ 0.01; ***p ≤ 0.001; ****p ≤ 0.0001). Tests, two-tailed unpaired Student’s t-test if not stated otherwise, and specific p-values used are indicated in the figure or legends.

### Viral IRESes drive translation of purified synthetic circRNAs in transfected cells

For in-cell assays, we transiently transfected purified circRNAs into human HEK293T cells and harvested them 24 hours post transfection (hpt). We hypothesized that Nluc reporters provide a more sensitive readout than EGFP. First, EGFP circRNAs were analyzed by FACS of EGFP+ cells and quantification of the measured EGFP signal (**Fig. 1E**). We titrated the amount of EGFP circRNA transfected (0.5, 1 and 2 μg) for robust EGFP+ cell detection. Using *hHBB* and HCV IRES as negative and positive controls, respectively, we find that 1 μg circRNA yields robust EGFP expression, with no further increase from 1 to 2 μg. These data suggested that the amount of EGFP circRNA that can be transfected and expressed saturated at 1 μg. The HCV IRES supported circRNA EGFP translation ∼3-fold compared to the *hHBB* control. We transiently co-transfected HEK293T cells with Nluc circRNAs and a linear capped *hHBB*-Fluc mRNA as a transfection control (**Fig. S1C**), and measured relative Nluc/Fluc signal from cellular lysates (**Fig. 1F**). *hHBB* showed lower activity than the empty construct, HCV high (3-fold), and CVB3 very high (1500-fold) IRES activity compared to empty, respectively, stemming from very efficient circRNA initiation (**Fig. S1D**). To control for the specificity of circRNA translation dependent on IRES-identity, we included the inverse sequence of CVB3 that reduced IRES-like activity to *hHBB* control level (**Fig. 1F**). Overall, these data indicated that both HCV and CVB3 IRESes can specifically recruit ribosomes to transfected synthetic Nluc circRNAs in human cells. Synthetic circRNAs are therefore a useful tool to study *bona fide* IRES functions, e.g. IRESes discovered in screens, in a system that is not affected by possible cellular processing artifacts. IRES activity in circRNAs also presents a benefit for long-term expression systems.

### IRES-mediated translation of synthetic circRNAs in *in vitro* translation systems

Synthetic circRNAs can also be useful for increased, upscaled recombinant protein expression in reductionistic cell-free *in vitro* translation extracts. We aimed to show that IRES-mediated circRNA translation is compatible even with more reductionistic systems that are better standardizable than in-cell transfection for sustained protein synthesis from long-lived circRNAs under controlled conditions. We applied Nluc circRNAs containing viral IRESes active in cells to the commonly used rabbit reticulocyte lysate (RRL) and wheat germ lysate (WGL) translation extracts, as well as to a recently developed human tripartite HEK293 cell-free translation system^82^ (**Fig. 2A**). Historically, cell-free systems have relied on actively translating models such as *Escherichia coli*, WGL^83^, insect cells, or RRL^84^. Although widely employed since the 1980s, RRL, and the generally translationally weaker WGL, do not always recapitulate cellular translation regulation^85^. We first compared RRL (**Fig. 2B**) and WGL (**Fig. 2C**) for IRES-dependent circRNA translation. Reporter translation was assessed by relative Nluc/Fluc luminescence and HA-Western blot analysis of Nluc protein levels using an Fluc mRNA as a reference. In RRL, HCV and CVB3 IRESes and CVB3 inverse sequence showed similar activity to the empty and *hHBB* controls (**Fig. 2B**). Consistent with previous reports, RRL did not distinguish between capped and uncapped mRNAs, indicating limited translational regulation^86^. We obtained similar results at 10-fold lower RNA concentrations, excluding effects of saturation or dynamic range, which suggested for example the lack of positive or the presence of negative ITAFs in RRL (**Fig. 2B**). Notably, high HCV and CVB3 activity as in in-cell assays was not observed. We tested the two IRESes in circRNAs in WGL and obtained very similar results, with high but unregulated translation across all constructs irrespective of IRES identity, seen by luciferase and by uniformly high Nluc signals by HA-specific Western blot (**Fig. 2C**). Thus, the IRESes tested appear inactive in both RRL and WGL.

**Figure 2.**
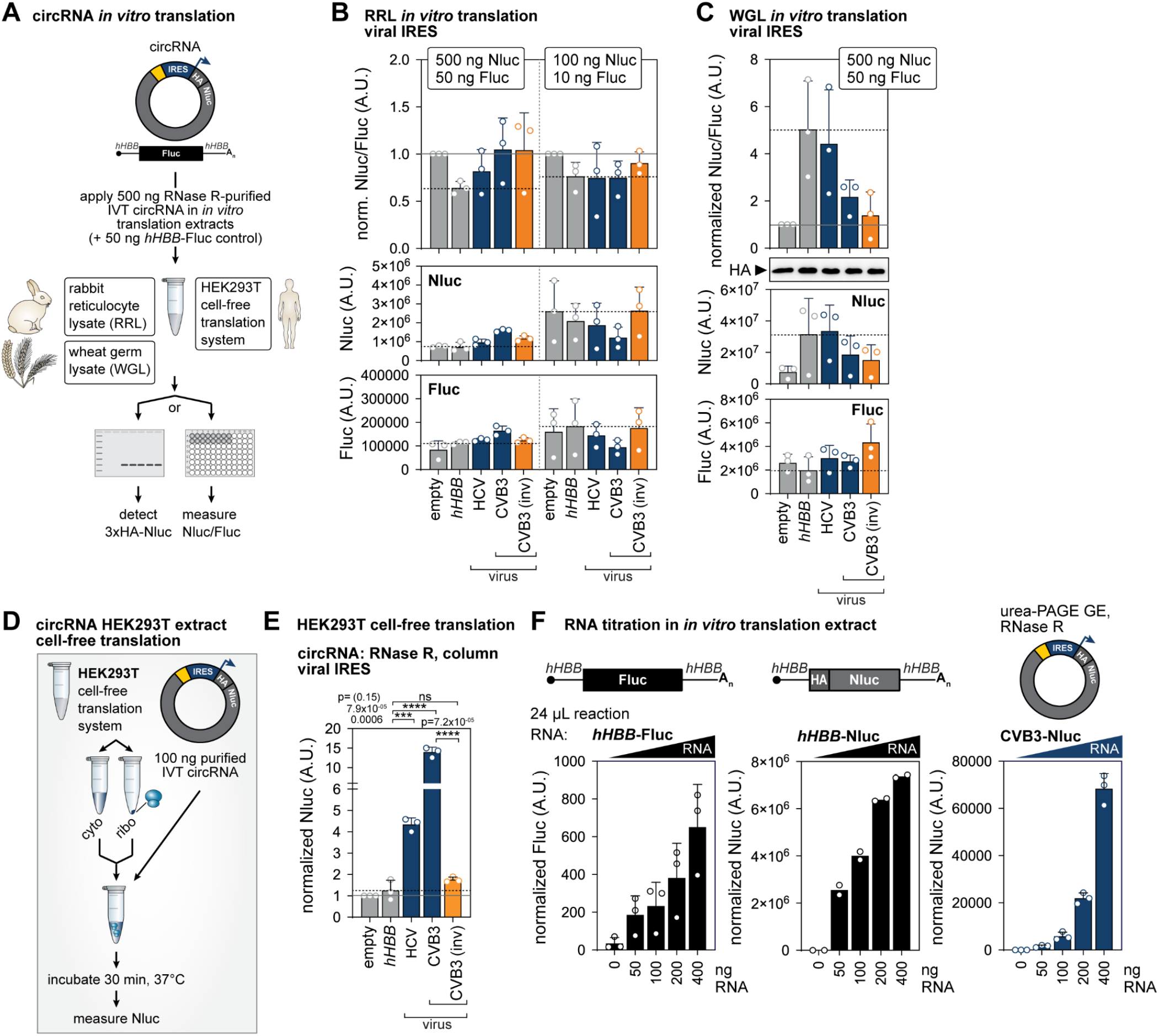
IRES-mediated circRNA-Nluc translation in cell-free *in vitro* translation across translation systems. **(A)** Experimental outline of the *in vitro* translation extracts used for IRES mediated translation of purified Nluc circRNAs. We evaluated the rabbit reticulocyte lysate (RRL), the wheat germ lysate (WGL), and the HEK293T cell-free *in vitro* translation systems.**(B)** Evaluation of translation of the different circRNAs in RRL including different tested viral IRES sequences. Nluc circRNA was co-translated with Fluc linear capped Fluc control RNA (ratio 1:10) with 500 ng Nluc circRNA (left) and 100 ng Nluc circRNA (right). CircRNA was purified by RNase R digestion and column purification. Bar graphs present the Nluc/Fluc ratio for each tested IRES sequence normalized to the empty vector control (top row), the raw Nluc data (second row), the raw Fluc data (third row); n=3.**(C)** Evaluation of translation of the different circRNAs in WGL including different tested viral IRES sequences. Nluc circRNA was co-translated with Fluc linear capped Fluc control RNA (ratio 1:10) with 500 ng Nluc circRNA. CircRNA was purified by RNase R digestion and column purification. Bar graphs present the Nluc/Fluc ratio for each tested IRES sequence normalized to the empty vector control (top row), the raw Nluc data (second row), the raw Fluc data (third row); n=3. HA-specific Western blot analysis shows respective Nluc protein levels.**(D)** Experimental outline of the HEK293T cell-free *in vitro* translation system used for IRES mediated translation of purified Nluc circRNAs.**(E)** Evaluation of translation of the different circRNAs in HEK293T extract including different tested viral IRES sequences. CircRNA was purified by RNase R digestion and column. Bar graphs present Nluc expression for each tested IRES sequence normalized to the empty vector control. Tested sequences are depicted in dark blue, inverse controls in orange, and controls in light grey; n=3.**(F)** Evaluation of translation in dependency of CVB3-circRNA concentration (right panel) and linear capped *hHBB*-Fluc mRNA (left), linear capped *hHBB*-Nluc mRNA (middle), and CVB3-Nluc circRNA (right), using a HEK293T *in vitro* translation extract (ReCet, Immagina). CVB3 containing circRNA was purified by urea-PAGE and RNase R digestion and column. Bar graphs present Nluc expression for each tested RNA concentration normalized to the no-RNA containing control reactions (background). Tested sequences are depicted in dark blue (circRNA) and black (linear mRNA). Reaction volumes were adjusted to 24 μL; n=3.

We next employed a newly improved HEK293-based translation system that separately purifies ribosomes, cytoplasm, and mRNAs, allowing controlled reconstitution and manipulation of the three components individually (e.g. by stress or drug treatment)^82^. This system enabled more physiological analysis of translation regulation, was applied to several cell types^82^, and is commercially available. In contrast, we established the human HEK293T-based tripartite cell-free system^82^ (**Fig. 2A, D**). We found that it recapitulates IRES-dependent regulation of Nluc circRNAs in cells (**Fig. 2E**). We showed high HCV and CVB3 IRES activity compared to *hHBB* (**Fig. 2E**), consistent with results from circRNA-transfected cells (**Fig. 1F**). The inverse CVB3 insert diminished activity, confirming specificity. We next examined whether circRNA input amounts in the reaction affected translation capacity. Excess RNA can lead to phosphorylation of eukaryotic initiation factor 2-alpha (eIF2α) which suppresses global cap-dependent translation initiation, as observed for mRNA titrations in the HEK293 extracts before^82^. It was unknown whether circRNAs are resistant to this effect due to lack of a cap. We added increasing amounts of either linear *hHBB*-Fluc mRNA or *hHBB*-Nluc mRNA^10^, or Nluc-circRNA with the strongest IRES (CVB3) to HEK293T translation extracts and monitored luminescence (**Fig. 2F**). Increasing amounts of linear mRNAs scaled linearly in translation but circRNA titrations scaled exponentially with higher overall translation. This enhanced circRNA translation may be due to more efficient loading of ribosomes onto circRNAs with more efficient ribosome recycling and re-initiation compared to linear mRNAs. Therefore, IRES-mediated initiation on circRNAs produced more protein in this system than cap-mediated initiation on linear capped mRNA. RRL and WGL are not suitable to study IRES-mediated circRNA translation due to their failure to support regulated internal initiation. However, the HEK293T cell-free translation system captures some features of translation regulation of certain IRESes in circRNAs.

### Select cellular IRESes can drive circRNA translation in cells and *in vitro*

Viral IRESes drive strong circRNA translation, but producing high protein levels over short periods of time may not be optimal for protein replacement and other therapies that require endogenous protein levels. We therefore selected six cellular IRESes previously shown to have IRES activity in plasmid-derived back-spliced circRNAs^81^ (**Fig. 3A**). Among picked cellular IRESes, IRES screen-derived *Chrdl1* and *Dlx1* have an ultraconserved 5′ UTR element^39^ and had the highest activity among cellular IRESes in plasmid-based circRNAs^81^. We also included the structured *Hoxa9* and *Hoxa5* 5′ UTR IRES-like translation enhancers, with previously assigned IRES activity in linear reporters^44,52,87,88^ and circRNAs^81^. We also included *c-Myc*^*38,89,90*^ and *Cofilin*^*91*^ IRESes, previously active in plasmid-based circRNAs^81^, in later experiments. We IVT-generated, RNase R- and column-purified these cellular IRES-encoding Nluc circRNAs (**Fig. 3B, Fig. S1A, B**), as for viral IRES circRNAs (**Fig. 1D**). IRES identity had little effect on circRNA generation and purification, except for *Dlx1*, where extended digestion (total 90 min) was needed to reach similar purity (**Fig. 3B**). We then transiently co-transfected HEK293T cells with Nluc circRNAs encoding cellular IRESes and a *hHBB*-Fluc mRNA, and measured relative Nluc/Fluc signals (**Fig. 3C**). Among cellular IRESes, *Hoxa5* and *Dlx1* were inactive, but the *Hoxa9* IRES-like element and *Chrdl1* were moderately active (2.8-fold and 1.5-fold more IRES activity than *hHBB*, respectively).

**Figure 3.**
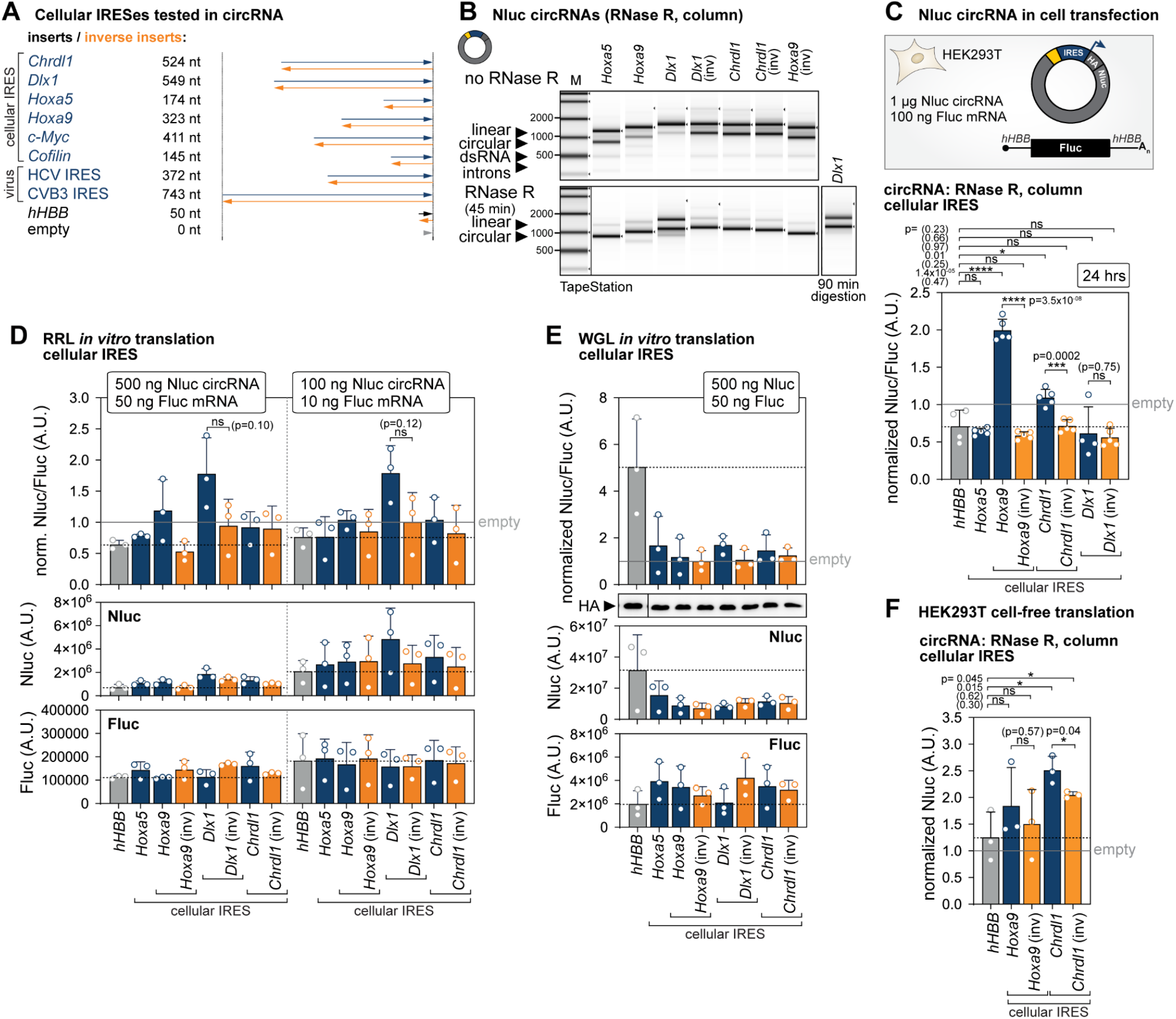
Cellular IRES-mediated circRNA-Nluc translation in cells and across cell-free *in vitro* translation systems. **(A)** Overview of the tested IRES sequences indicating length and viral or cellular origin of the IRESes, and their respective inverse controls.**(B)** Quality control, using the High Sensitivity RNA ScreenTape, of the generated Nluc circRNAs before purification (upper panel) and after RNase R digestion (lower panel). Remaining contaminants can be observed as light grey bands.**(C)** IRES activity of the tested sequences was evaluated 24 hpt using Nluc circRNAs as in **Fig 1F**. RNA was purified by RNase R digestion and column. Tested sequences are depicted in dark blue, inverse controls in orange. 1 μg of circRNA was co-transfected with 100 ng *hHBB*-Fluc linear capped control RNA. The same *hHBB* data as in **Fig. 1F** is included in both viral and cellular panels. Nluc data were normalized to Fluc transfection control and Nluc/Fluc ratios are normalized to empty control; mean ± SD, n=3-5.**(D)** Evaluation of translation of the different circRNAs in RRL including different tested IRES sequences. Nluc circRNA was co-translated with Fluc linear capped Fluc control RNA (ratio 1:10) with 500 ng Nluc circRNA (left) and 100 ng Nluc circRNA (right). CircRNA was purified by RNase R digestion and column purification. The same *hHBB* data as in **Fig. 2B** is included in both viral and cellular panels. Bar graphs present the Nluc/Fluc ratio for each tested IRES sequence normalized to the empty vector control (top row), the raw Nluc data (second row), the raw Fluc data (third row); n=3.**(E)** Evaluation of translation of the different circRNAs in WGL including different tested IRES sequences. Nluc circRNA was co-translated with Fluc linear capped Fluc control RNA (ratio 1:10) with 500 ng Nluc circRNA. CircRNA was purified by RNase R digestion and column purification. The same *hHBB* data as in **Fig. 2C** is included in both viral and cellular panels. Bar graphs present the Nluc/Fluc ratio for each tested IRES sequence normalized to the empty vector control (top row), the raw Nluc data (second row), the raw Fluc data (third row); n=3. HA-specific Western blot analysis shows respective Nluc protein levels.**(F)** Evaluation of translation of the different circRNAs in HEK293T extract including different tested IRES sequences. CircRNA was purified by RNase R digestion and column. The same *hHBB* data as in **Fig. 2E** is included in both viral and cellular panels. Bar graphs present Nluc expression for each tested IRES sequence normalized to the empty vector control. Tested sequences are depicted in dark blue, inverse controls in orange, and controls in light grey; n=3.

As all four cellular IRESes were active at 24 hpt in plasmid-based circRNA systems in the same cell type^81^, these differences in activity in synthetic circRNAs may indicate IRES requirements for “nuclear experience”^92^, including RNA modifications such as m^6^A^33^ and ITAFs, whose regulatory contribution was not reproduced with exogenous circRNAs. Previously, *Dlx1* was twice as active as *Chrdl1* in plasmid-derived circRNAs^81^, which was reversed in *in vitro* circRNAs. *Hoxa9* was comparable in activity to *Chrdl1*, and here about twice as active. For *Chrdl1* and *Dlx1*, IRES activity is known to vary across different cell types^39^, but it is unknown for *Hoxa5*. circRNAs applied across cell types may reflect factor requirements due to differential expression of ITAFs or specific RNA modifications, but are also a great way to filter out false positive IRES activity possibly detected by older, orthogonal methods^47,93^. However, absence of IRES activity in circRNAs, particularly in synthetic circRNAs, does not exclude IRES activity in other systems and cell types. The inverse sequences of *Hoxa9, Chrdl1*, and *Dlx1* all reduced IRES-like activity to *hHBB* (**Fig. 3C**). Overall, these data indicate that 2/4 cellular IRESes tested (*Hoxa9, Chrdl1*) can specifically recruit ribosomes to transfected synthetic Nluc circRNAs in human cells.

We next tested *Hoxa5, Hoxa9, Chrdl1*, and *Dlx1* for their activity in *in vitro* translation systems. When compared in RRL (**Fig. 3D**) and WGL (**Fig. 3E**), cellular IRES-dependent circRNA translation compared to a Fluc mRNA reference, yielded similar results than for viral IRESes (**Fig. 2B**). In RRL, most IRESes and their inverse sequences showed similar activity to the empty and *hHBB* controls, and their circRNAs were translated irrespective of the IRES (**Fig. 3D**). Only *Dlx1* showed a trend towards increased activity, however non-significant, that was diminished by its inverse insert. However, only the *Dlx1* circRNA required extended RNase R digestion to enrich circRNAs suggesting that residual linear uncapped RNA may contribute to translation. As observed for viral IRESes, we tested the four cellular IRESes in circRNAs in WGL and obtained high but unregulated luciferase translation and uniformly high Nluc signals by HA-specific Western blot across all constructs unrelated to IRES identity (**Fig. 3E**). Thus, both RRL and WGL did not mirror translation regulation by cellular IRESes in circRNAs. In contrast, in the human HEK293T-based tripartite cell-free system^82^ recapitulated cellular IRES-dependent regulation of Nluc circRNAs for *Chrdl1* (**Fig. 3F**), consistent with results from circRNA-transfected cells (**Fig. 3C**). The inverse *Chrdl1* insert reduced activity, confirming specificity. However, the *Hoxa9* IRES-like element compared to its inverse control was not active in this system (**Fig. 3F**), unlike in cells (**Fig. 3C**). These results highlighted the differences between in-cell and *in vitro* settings and suggested that regulatory mechanisms on cellular IRESes are not fully recapitulated in translation extracts.

### Comparison to plasmid-based back-spliced and ribozyme-mediated circRNA systems

If nuclear history plays a role in IRES activity, generating circRNA in the nucleus of transfected cells may affect the results. After observing that IRES identity matters for translation activity in synthetic circRNAs across in-cell and *in vitro* systems, we aimed to compare all IRESes to their activity in plasmid-derived circRNAs. Generating circRNAs intracellularly upon plasmid transfection is more broadly used but provides less control with regards to RNA processing and the final mature circRNA generated compared to synthetic synthesis and RNA transfection. For this, we compared *in vitro* generated circRNAs to two commonly used plasmid transfection-based systems that rely on either nuclear back-splicing or ribozyme-mediated cleavage and ligation for circularization. We previously used a split-EGFP reporter plasmid with permuted ZKSCAN1 introns, where the introns mediate nuclear back-splicing to fuse two EGFP fragments in a pre-mRNA into a full-length (FL) EGFP ORF^12,29,76,81^ (**Fig. S2A**). The IRES is located upstream of the 5′ EGFP portion. This approach has enabled IRES-screening and -mutagenesis^12,81^, and was important for the optimization of large gene delivery^94^. All viral and cellular IRESes we tested here supported IRES-mediated translation in this system^81^. It is suitable for studying endogenous IRES-based translation regulation as nuclear-generated circRNAs mimic physiological mRNA generation and cytoplasmic export. However, the ZKSCAN1 intron-containing reporter pre-mRNA is often inefficiently back-spliced. Previous reports suggest that back-spliced reporters may also undergo *trans*-splicing that generates long linear capped mRNAs encoding FL EGFP in a tandem reporter mRNA that may contribute to overall fluorescence^95–97^ (**Fig. S2B**). To assess if and to which extent *trans*-splicing occurs in the ZKSCAN1 intron system, we chose a viral (CVB3) and cellular (*Hoxa9*) IRES (**Fig. S2C**). We sorted EGFP+ cells at 5 dpt, and subjected total RNA to RNase R treatment before Northern blot analysis. The probe used is specific for the EGFP junction region that should capture the circRNA and potential *trans*-spliced RNA species if expressed (**Fig. S2B**). For both CVB3 and *Hoxa9* IRESes, the circRNA was clearly the dominant species, before and after RNase R, with no major linear RNAs detected (**Fig. S2D**). Thus, the ZKSCAN1 intron-based plasmid system primarily generates circRNAs in HEK293T cells.

A second plasmid transfection-based system is the recently optimized Tornado (<u>T</u>wister-<u>o</u>ptimized RNA for <u>d</u>urable <u>o</u>verexpression) translation system^95,98^. It does not require nuclear splicing but twister ribozyme cleavage^99^ and RtcB ligase-mediated ligation. It uses a split Nluc reporter and is thought to be superior to back-spliced systems for efficient cytoplasmic mRNA circularization and expression^95^ (**Fig. S3A**), with minimal linear precursors. The developers of the Tornado system found the viral CVB3, HRV-B3, HCV and EMCV IRESes to drive efficient circRNA translation^95^. However, four cellular IRESes (*LIMA1, CEP50, TGFBRG, CHD2*) with high translational activity in an IRES screen in circRNAs^12^ had low activity in the Tornado system^95^, none included here. We explored this further and tested seven diverse cellular IRESes and their inverse controls (*Chrdl1, Dlx1, Bcl2, c-myc, Cofilin, Hoxa5, Hoxa9*), all previously active IRESes in the ZKSCAN1 system^81^, compared to viral CVB3 and HCV IRESes, and the *hHBB* negative control (**Fig. S3B**). At 4 dpt, we observed high activity for CVB3 and HCV, but all seven cellular IRESes failed to promote translation in this system (**Fig. S3B**). We assumed that either remaining structured ribozyme regions interfere with IRES-ribosome recruitment, or required IRES-interacting RBPs as ITAFs were missing which was not compatible with the Tornado translation system.

### Immunogenicity of diverse IRESes in synthetic circRNAs

Relevant for therapeutic application of synthetic circRNAs, we next tested if IRES identity affects circRNA immunogenicity^25,29,69–72^. We asked whether the viral or cellular origin of an IRES or the status of high or poor translation initiation contributes to circRNA immunogenicity. To assign immune sensing specifically to IRES inserts, we generated highly purified circRNAs free of immune-stimulatory linear IVT side products. RNase R treatment alone is not sufficient to obtain highly pure circRNA^29^ as it retains, besides circRNAs, also other RNase R-resistant species including structured linear RNAs and short dsRNA fragments^25^ (**Fig. 1A**). Even trace amounts of linear RNA contaminants, present even after HPLC purification, were shown to trigger an immune response^25^. We therefore purified circRNAs by 4% urea-PAGE gel extraction (GE) and RNase R-treated the excised circRNA after. We confirmed that circRNAs were free of IVT side products and introns by TapeStation capillary analysis, exemplified for the *Hoxa9* IRES-like element (**Fig. 4A**). We noted that on urea-PAGE gels, the circRNA band ran higher than the linear product, obvious from RNase R treatment, but this migration pattern inverted on TapeStation RNA chips with linear RNA running higher (**Fig. 4A**). This procedure yielded comparable highly pure circRNAs across all nine constructs including IRESes HCV, CVB3, *Hoxa9, Chrdl1*, as well as *Coflin* and *c-Myc*, and their inverse sequences (**Fig. 4B, S4**). We then co-transfected the gel-extracted and RNase R-digested Nluc circRNAs containing the HCV, CVB3, *Hoxa9, Chrdl1, Coflin* and *c-Myc* IRESes into HEK293T cells and found comparable relative IRES activities as with RNase R-only circRNAs for the viral IRESes (**Fig. 4C**, see also **Fig. 1F, 3C**). Viral HCV and CVB3 IRES were highly active, with HCV being more active in purified circRNAs compared to RNase R-enriched only (**Fig. 1F**). The *Hoxa9* IRES-like element did not confirm IRES activity in purified circRNAs after GE over its inverse control. *Cofilin* and *c-Myc* were 3- and 2-fold more active than *hHBB* (**Fig. 4C**). Thus, differences in circRNA preparation purity affected select IRES-specific activity for circRNA translation.

**Figure 4.**
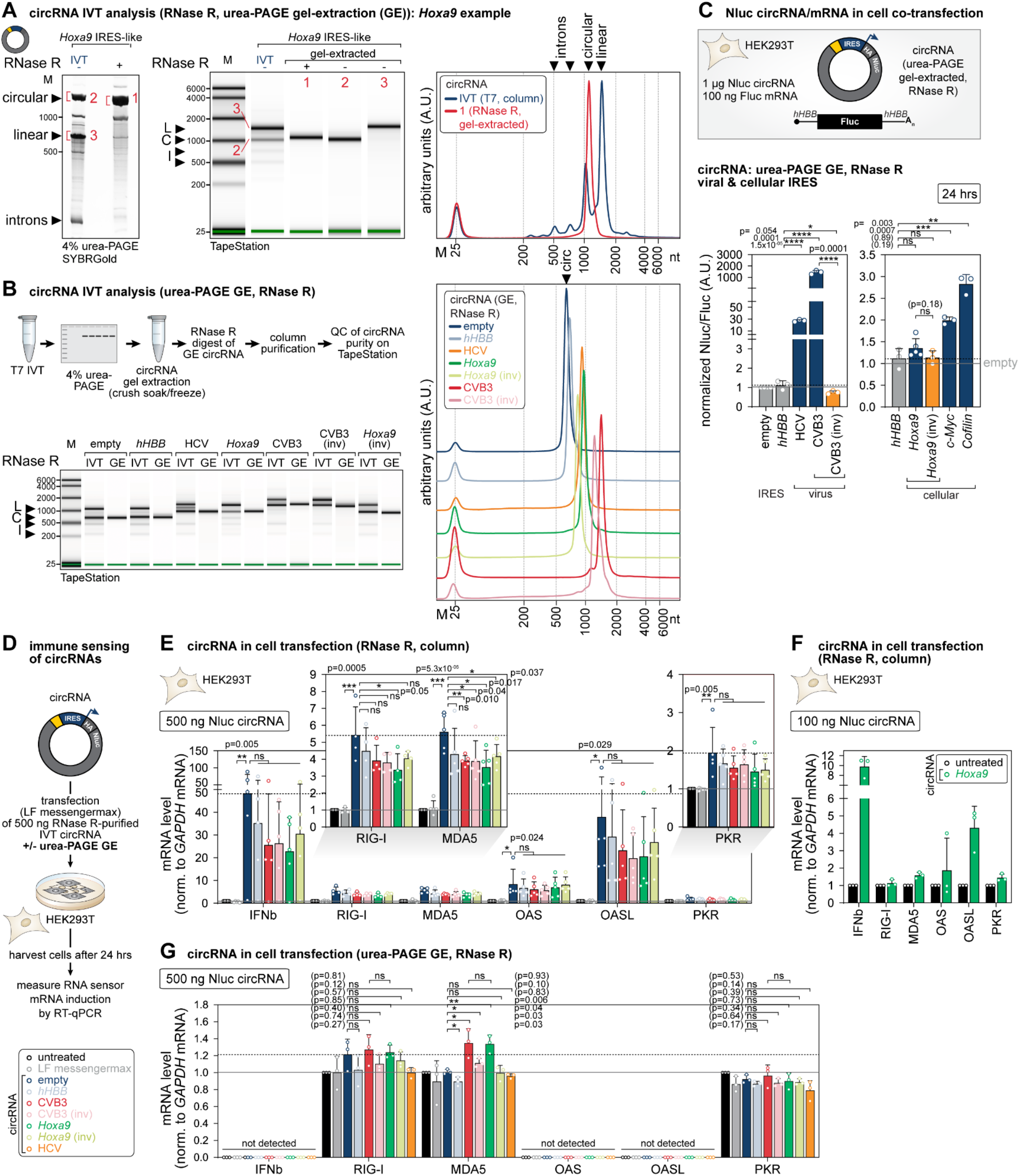
circRNA sensing and evaluation of IRES-specific immunogenicity of urea-PAGE gel extracted circRNAs of high purity. **(A)** Optimized circRNA purification after IVT, exemplified for the *Hoxa9* IRES-like insert. 4% urea PAGE-gel after SYBR Gold staining (left). IVT RNA is shown in lane 1, RNase R digested IVT RNA is shown in lane 2. The remaining upper band (1) is formed by the circular RNA as well as the upper band of the untreated IVT RNA (2). Linear non-circularized RNA species are forming band (3). Circular and linear RNA were gel-extracted (GE) (band (1) - (3)) and analyzed using the Agilent RNA high sensitivity screen tape (middle). The corresponding electrogram data of the unpurified IVT RNA (dark blue) as well as the RNase R and gel-purified circRNA (red) are shown (right). Intron sequences, linear and circular RNA species are indicated according to size.**(B)** Visualization of the circRNA purification pipeline combining urea-PAGE gel extraction (GE) and RNase R digestion (upper panel). Quality control of the obtained circRNA using the Agilent RNA high sensitivity screen tape (lower panel). Shown are the unpurified IVTs next to the RNase R and gel-purified circRNAs. The insert sequences are indicated above. Linear RNA-containing band (L), circRNA-containing band (C) and intron band (I) are indicated. Corresponding electrogram data are shown (right; colored according to the tested IRES insert).**(C)** IRES activity of the tested sequences was evaluated 24 hpt in Nluc circRNA as in **Fig. 1F**. RNA was purified by urea-PAGE, RNase R digestion and column. IRES sequences are displayed in dark blue, inverse controls in orange. Activities of IRESes of viral and cellular origin are depicted separately, with the same *hHBB* data included in both viral and cellular panels. 1 μg of circRNA was co-transfected with 100 ng *hHBB*-Fluc linear capped control mRNA. Nluc data were normalized to Fluc transfection control and ratios are normalized to empty vector control; n=3-5.**(D)** Experimental outline of the immune sensing experiment evaluating the immunogenicity of Nluc circRNAs and containing different IRES sequences (indicated by color, lower panel) after different purification methods. RNA was purified by urea-PAGE, RNase R digestion and column purification or RNase R only and column purification. HEK293T cells were harvested 24 hpt and RNA sensing was evaluated by RT-qPCR quantification of immune sensor RNA expression normalized to human GAPDH mRNA levels.**(E)** Evaluation of purified circRNA immunogenicity dependent on encoded IRES sequences 24 hpt. Transfected circRNA was RNase R digested and column purified. mRNA expression levels of different immune sensors were quantified (normalized to human GAPDH mRNA levels) after 500 ng circRNA transfection. Colors of the bar graphs according to the tested IRES sequences. mRNA expression levels are shown relative to untreated cells; n=4.**(F)** Same experiment as in (E) using 100 ng circRNA transfection into HEK293T cells, exemplary for the *Hoxa9* IRES-like element. mRNA expression levels are shown relative to untreated cells; n=3. See also **Fig. S5.(G)** Evaluation of purified circRNA immunogenicity dependent on IRES sequence 24 hpt. Transfected circRNA was urea-PAGE extracted, RNase R digested, and column purified. mRNA expression levels of different immune sensors were quantified (normalized to human GAPDH mRNA levels) after 500 ng circRNA transfection. Colors of the bar graphs according to the tested IRES sequences. mRNA expression levels are shown relative to untreated cells. Ct values of “not detected” sensors were too high for reliable quantification; n=3.

Circularization as such does not alter overall RNA structure compared to linear mRNA, as reported for the EMCV IRES^29^. We therefore asked whether long dsRNA regions within IRES structures from viral or cellular origin contribute to innate immune activation. dsRNA elements in circRNAs may sequester and inhibit PKR^70,72^. Whether IRESes of different origin are sensed differently by RNA sensors in exogenous circRNA is unclear. To assess this, we transfected circRNAs purified either by RNase R and column, or gel-extracted, into HEK293T cells and measured their immune stimulatory effect by mRNA induction of five RNA sensors (RIG-I, MDA5, 2′-5′-oligoadentylate synthase 1 (*OAS1*), OAS-like protein (*OASL*), PKR) by RT-qPCR (**Fig. 4D**). Compared to untreated cells or transfection reagent only, six RNase R-only purified circRNAs (empty, *hHBB*, CVB, CVB (inv), *Hoxa9* and *Hoxa9* (inv)) all induced comparable relative levels of all five sensors regardless of IRES identity (**Fig. 4E**). Type I interferons such as interferon beta (IFN-β) are produced by various cell types in response to viral infection, also by non-professional immune cells^75,100^. IFN-β mRNA induction here served as a marker for cytosolic RNA sensing and antiviral host immune signaling, which was similar across all circRNAs tested. There was no difference in sensor mRNA induction between empty and *hHBB* inserts compared to actively translated IRESes or inverse controls. Thus, IRES identity and circRNA translation state seemed not to determine the level of sensing. Subtle IRES-specific effects may be invisible due to strong sensing of circRNA IVT contaminants. Indeed, this was supported by the finding that the sensing effect scaled with the amount of transfected RNase R-only purified circRNAs, or in other words with the level of impurities in the RNA sample: 100 ng circRNA globally induces the sensors less than 500 ng (**Fig. 4F**; **Fig. S5**). We next transfected highly purified (gel-extracted and RNase R-treated) circRNAs. Very clearly, they globally reduce or completely abolish immune activation (**Fig. 4G**). The level of RNA sensor mRNA induction did not seem to depend on IRES identity and translation state. CVB3 was sensed a bit more than CVB (inv) by MDA5, but there was no difference in MDA5 induction by CVB3 or *Hoxa9*. We concluded that IRES identity and translation state did not influence immunogenicity as assessed by sensor mRNA induction in HEK293T cells. The responsible RNA sensing pathways need to be investigated in immune cells to clearly delineate IRES-dependent immunogenicity. These data suggested that neither viral and cellular IRES origin, nor cell-foreign *td* introns, are the primary determinant for sensing of circRNAs as foreign. Rather, the purification status and presence of IVT contaminants are predominantly responsible for immune recognition of circRNAs.

### Engineering of HCV IRES-mediated circRNA translation with rational design mutations

Rational engineering of IRESes in synthetic circRNAs may allow for fine-tuning of protein expression and enable functional IRES characterization. For this, we chose the HCV IRES that mediated robust IRES activity in transfected cells (**Fig. 1F, 4C**) and in cell-free HEK293T extracts (**Fig. 2E**). From cryo-EM reconstructions of HCV IRES-ribosome complexes, we know the interactions of the HCV IRES with eIFs and the ribosome to recruit the 40S ribosomal subunit via eIF3 and rRNA contacts^101,102^. We made use of previously isolated mutants in the HCV IRES that delete the dII domain (ΔdII) and target the interaction site of 18S rRNA expansion segment 7 (ES7S), a structural anchor for the HCV IRES (dIIId GGG-to-CCC), that both diminish HCV IRES translation initiation activity in in-cell assays^103,104^ (**Fig. 5A**). We found that in transfected cells, the activity of the HCV IRES to drive circRNA translation was highly reduced to a similar extent with both ΔdII and dIIId GGG-to-CCC HCV mutants (**Fig. 5B**). Thus, the effects of inhibitory HCV IRES mutants previously seen in linear or bicistronic reporter mRNAs can be fully recapitulated in circRNA reporters. We next asked whether we can see the same effect of these mutants in the cell-free HEK293T extract (**Fig. 5C**). When we employed HCV mutant circRNAs to these extracts, we saw an overall lower activity of the WT HCV IRES of 2.7-fold (**Fig. 5C**) compared to 22-fold in cells (**Fig. 5B**) over the empty control. The dIIId GGG-to-CCC mutant specifically reduced translation to 2-fold over the control, but the ΔdII mutant was equally active than the HCV IRES (**Fig. 5C**). These data indicated that the cell-free translation system cannot fully recapitulate translation regulation as in cells and requires further optimization to allow characterization and engineering of dynamic translation kinetics. However, targeted mutagenesis of IRESes to engineer IRES-dependent translation seen in linear RNAs can be fully reconstituted with circRNA reporters.

**Figure 5.**
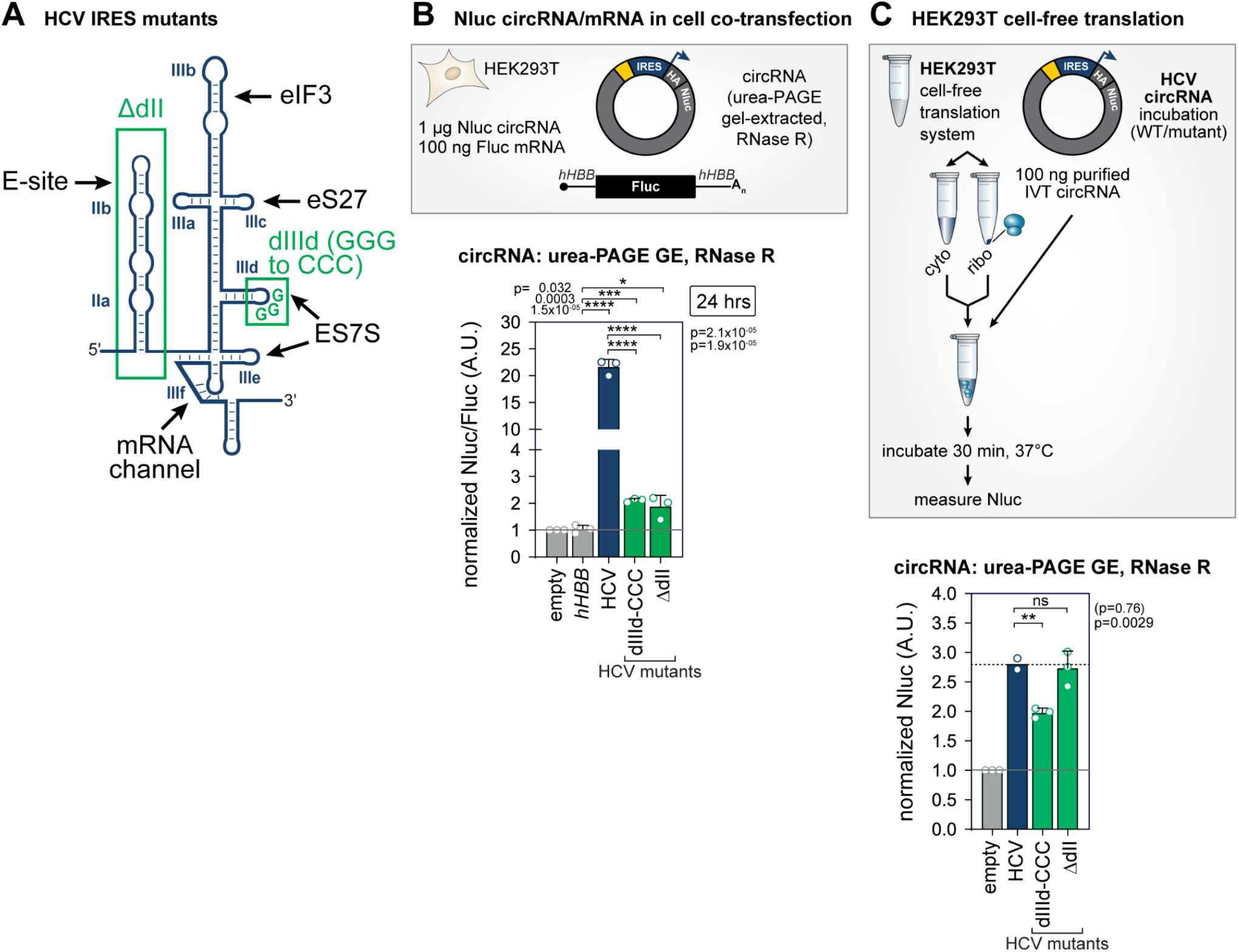
Engineering of HCV IRES-mediated circRNA-Nluc translation in cell-free translation systems. **(A)** Schematic illustration of the viral HCV IRES structure. For ribosome recruitment essential binding sides are indicated with black arrows, tested mutations which should impact the IRES function are depicted in green.**(B)** Evaluation of circRNA translation 24 hpt. circRNAs include WT HCV IRES (dark blue), HCV mutants (green) and the *hHBB* negative control (grey). CircRNAs were urea-PAGE and RNase R and column purified. Linear capped *hHBB*-Fluc control mRNA was co-transfected (ratio 1:10). Nluc values are normalized to Fluc transfection control. Nluc/Fluc ratios are normalized to the empty vector control; n=3.**(C)** Experimental outline of the HEK293T cell-free *in vitro* translation system used for HCV IRES-mediated translation of purified Nluc circRNAs. Evaluation of circRNA translation dependent on the HCV IRES. circRNAs include WT HCV IRES (dark blue), HCV mutants (green) and empty negative control (grey). CircRNAs were urea-PAGE and RNase R and column purified. Nluc values are normalized to the empty vector control; n=2-3.

## DISCUSSION

We present a comprehensive comparison and optimization of methods to investigate diverse IRES RNA function in synthetic circRNA reporters in cells and *in vitro*. How IRESes mediate circRNA translation, or if IRESes in circular and linear mRNAs act differently, is understudied. We find viral IRESes (**Fig. 1, 2**) to be generally more potent in circRNAs than the tested cellular IRESes (**Fig. 3**). This reflects the biology of naturally occurring mRNA elements that contribute to translation, particularly the maintenance of cap-independent translation in response to stress. This is orders of magnitude lower than the strong viral translation that occurs during infection. Low but sustained translation from cellular IRESes may be advantageous for long-term protein replacement therapy in contrast to high and transient antigen expression from short-lived mRNA vaccines. Cell-type specificity of IRES activity in circRNA therapeutics may further enable customizable tissue-targeted therapies. Many viral IRESes sustain host-dependent translation of viral mRNAs upon host infection, often assisted by required ITAF protein cofactors^36,37^. Such specificity may stem from tissue-specific ITAF availability or ribosome-associated factors. The selected cellular IRESes have specific expression patterns. Among cellular IRESes, *Chrdl1* and *Dlx1* were identified as ultraconserved 5′ UTR elements in a screen for cell type-specific IRES-mediated translation^39^, and ranked among the highest active cellular IRESes in plasmid-based circRNAs^81^. These IRESes are important for mouse embryonic development and mRNA translation efficiency^39^. Dlx1 acts as a Hox transcription factor in craniofacial patterning and neuronal differentiation and survival^105,10639^. We also include the structured *Hoxa9* and *Hoxa5* 5′ UTR IRES-like translation enhancers from the mouse HoxA gene cluster, with previously assigned IRES activity in linear reporters^44,52,87,88^ and circRNAs^81^. *Hoxa* IRES-like elements regulate IRES-dependent, ribosome-directed translation in mouse embryo anterior-posterior patterning of the axial skeleton^52,87^. The *Hoxa9* IRES-like element recruits 40S ribosomal subunits for initiation through mRNA–rRNA interactions with rRNA ES9S^44^. We also include *c-Myc*^*38,89,90*^ and *Cofilin*^*91*^ IRESes. As viral IRESes (HCV, CVB3) are consistently active across circRNA assays, only a subset of the tested cellular IRESes (*Hoxa9, Chrdl1, Cofilin, c-Myc*) function in synthetic circRNA reporters (**Fig. 3, 4**). However, all were previously active in the ZKSCAN1-based circRNA plasmid system in the same cell type^81^. This suggests that IRESes usually embedded in endogenous 5′ UTRs in linear mRNAs may display stronger requirements for cofactors, “nuclear experience” of RBP remodeling, in-cell RNA processing, or ITAF-based structure stabilization for efficient ribosome recruitment. This discrepancy of cellular IRES performance in plasmid- and back-splicing-derived compared to synthetic circRNAs indicates that these layers of RNP remodeling cannot be fully recapitulated with synthetic circRNAs containing certain cellular IRESes. Different IRES activities across systems may also be explained by certain IRESes adopting distinct, more or less ribosome-competent RNA conformations in linear mRNAs and circRNAs that affect the dynamics and initiation efficiency. Integrating this knowledge will help design customized IRESes in circRNA therapeutics with the potential to be cell type-specific, more efficient, and useful for longer-lasting treatments than achievable with linear mRNAs.

Synthetic circRNAs offer a platform to dissect the biology of IRESes and translation enhancers, and to reveal rules of cap-independent translation by functional RNAs and m^6^A-mediated initiation by eIF3 and ribosome recruitment^107^. Cellular IRESes may be further m^6^A-modified to enhance ribosome binding. Synthetic circRNAs allow distinguishing RNA structure- and sequence-based IRES activity from m^6^A modification-driven initiation. Even clearer than plasmid-based back-spliced circRNA systems, synthetic circRNAs can robustly filter out false positive IRES activity. However, lack of activity in circRNAs does not exclude IRES function in other contexts. We find cellular IRESes to be globally inactive in the Tornado plasmid-based translation system (**Fig. S3**), which may be caused by inhibitory intermolecular interactions between ribozyme remnants and IRES inserts in *cis* that do not interfere with robust viral IRES function. Indeed, it was recently shown that extensive interactions between several viral IRESes and certain cargo sequences in circRNAs compromise IRES structural integrity, thereby impairing circRNA translation^108^.

Regarding immunogenicity, there is no consensus which RNA sensors are activated by circRNAs. Sensing of circRNAs is thought to depend on self or non-self intron remnants^71^. We focused on IRES identity and viral versus cellular origin that may be sensed differently in circRNAs due to different RNA structure, long dsRNA regions, or factor recruitment across IRESes. We find that circRNA sensing is primarily determined by RNA preparation purity rather than IRES identity. A more detailed answer will require RNA sensing readouts in immune cells that preserve the full receptor repertoire and sensing pathways. In our experiments, induction of IFN-β mRNA after circRNA transfection reflects the activation of cytosolic RNA sensing pathways rather than circRNA delivery or translation. It is typically interpreted as evidence of immunogenic RNA or contaminants in the circRNA preparation (immune-stimulatory dsRNA IVT contaminants, residual 5′-triphosphate linear RNA, aberrant structures in circRNA and IRESes, and/or incomplete RNase R purification) (**Fig. 4**). We can use our assay to find “immune-silent” IRESes in circRNA designs, as long as the circRNA is urea-PAGE extracted and RNase R treated. Purified circRNA with active IRESes minimizes activation of MDA5, but abolishes RIG-I or PKR induction (**Fig. 4**). Indeed, circRNAs do not contain the 5′-triphosphate motif canonically required for RIG-I activation. RIG-I was suggested to transiently interact with circRNAs devoid of host nuclear proteins, leading to a typical RIG-I-mediated antiviral response^29^. It has also been found that circRNA synthesized by T4 RNA ligase without extraneous fragments exhibit minimized immunogenicity^72^. Cells distinguish between self (e.g. ZSCAN1 intron) and non-self (e.g. *td* intron) circRNAs based on the introns from which they were produced, perhaps because mature human circRNAs are associated with diverse RNA-binding proteins^29^. circRNAs recognized as foreign activating immune signaling raises the question of circRNA involvement in autoimmune diseases. Conversely, exogenous circRNAs may boost immune activation for therapeutic purposes. circRNAs are thought to be intrinsically less immunogenic than linear mRNAs^25,109^. However, whether incorporation of N1-methyl-pseudouridine (m^1^Ψ), as is necessary in linear vaccine mRNAs to reduce immune sensing^1–5^, is also important to be incorporated in circRNAs, has, to our knowledge, not been broadly discussed or studied.

We conclude that synthetic circRNA translation is constrained by ITAF dependence and RNA processing history of the circRNA itself, critical for cell type-specific RNA therapeutic applications. Our studies show that *in vitro* synthesized circRNAs, particularly sensitive Nluc reporters, provide an elegant system to investigate IRES-mediated translation from viral and cellular IRES origin in transfectable cells or cell-free systems. Recent cell-free systems are better at recapitulating physiological regulation but require further optimization. For synthetic biology, *in vitro* circRNAs are beneficial as they do not rely on the cellular splicing machinery, and allow for rigorous quality control before transfection or upscaled recombinant protein synthesis. We establish robust protocols for both in-cell and *in vitro* translation assays and clearly confirm specific IRES activity from several viral and cellular IRES elements in them. These findings highlight circRNAs as a reliable system to study IRES-mediated translation and are directly relevant for the development of emerging synthetic mRNA medicine.

## DATA AVAILABILITY

### Lead Contact

Further information and requests for resources and reagents should be directed to and will be fulfilled by the Lead Contact, Kathrin Leppek.

### Materials Availability

All plasmids generated in this study are available upon request and will be fulfilled by the Lead Contact, Kathrin Leppek.

### Data Availability

The source data of this paper are collected in the following database record: X.

Expanded view data, supplementary information, appendices are available for this paper at X.

## AUTHOR CONTRIBUTION

P.K. and K.L. conceived, and K.L. supervised the project; P.K. and K.L. designed the experiments, and P.K. performed experiments and analyzed the data; F.S.W.A. and A.H.L. provided HEK293T cell-free translation extracts and technical support in experimental design; C.L. and C.-K.C. performed ZKSCAN1-specific Northern blot experiments; C.-K.C. provided technical support in circRNA IVT and purification strategy; S.N. provided the *hHBB*-Fluc reference mRNA and performed RT-qPCR experiments; A.A. and M.C. (Immagina) generated and provided HEK293T cell-free translation kits; P.K. performed the rest of the experiments and analyses. K.L. prepared all figures. K.L. and P.K. wrote the manuscript with input from all co-authors.

## ACKNOWLEDGEMENT

We would like to thank the Leppek lab members for support and constructive criticism of the work, and Lisa Nauroth for technical assistance in the Leppek lab. We thank Maria Barna, Ina Huppertz, and Ranen Aviner for critical reading of the manuscript. We thank Maria Barna (Stanford University, Stanford, CA 94305, USA) for providing RNase R reagents, Howard Y. Chang (Stanford University, Stanford, CA 94305, USA; Amgen Research, South San Francisco, CA 94080, USA) for providing plasmid mRuby-circEGFP-ScaI-v3 generated by C.-K. Chen (Washington University School of Medicine, St. Louis, MO 63110, USA) as the basis for generating plasmids in this study; and both Howard and Maria for valuable discussions on the manuscript. We thank Katrin Reiners (Institute of Clinical Chemistry and Clinical Pharmacology, University Hospital Bonn, University of Bonn) for technical advice on the Attune FACS. Work in the Lund lab was supported by grants from The Novo Nordisk Foundation (NNF18OC0030656 and NNF21OC0071919) and the Independent Research Fund Denmark (10.46540/4262-00065B). Work in the Chen lab was funded by NIH (R00HG011475) to C.-K.C.. Work in the Leppek lab was supported by a Human Frontier Science Program (HFSP) Early Career Award (RGEC32/2023) to K.L., Fritz Thyssen Foundation project funding (10.25.1.026MN) (S.N.) to K.L., and a Ph.D. position funded by the Strengthening the Equal Opportunity Process (STEP) Program of the University of Bonn (S.N.) to K.L.. K.L. is supported by the Cluster of Excellence ImmunoSensation3 funded by the Deutsche Forschungsgemeinschaft (DFG, German Research Foundation) under Germany’s Excellence Strategy – EXC2151 – 390873048, and start-up funds of the Medical Faculty at the University Hospital Bonn and University of Bonn, Germany.

## DECLARATION OF INTERESTS

K.L. and Maria Barna (M.B.) are inventors on patents related to the *Hoxa9* P4 stem-loop and RNA therapeutics and their various uses. P.K., M.B. and K.L. are inventors on a patent related to IRES-like elements in plasmid-based and synthetic circRNA reporters and RNA therapeutics and their various uses. M.C. is Chief Executive Officer (CEO) and shareholder of IMMAGINA Biotechnology S.r.l.; A.N. is an employee of IMMAGINA Biotechnology S.r.l;. “ReCet” is a proprietary technology commercialized by IMMAGINA Biotechnology S.r.l.; F.S.W.A. and A.H.L. serve as scientific advisors and members of the Scientific Advisory Board of IMMAGINA Biotechnology S.r.l.

### Declaration of generative AI and AI-assisted technologies in the manuscript preparation process

During the preparation of this work the author(s) used ChatGTP5.3 in order to improve grammar. After using this tool/service, the author(s) reviewed and edited the content as needed and take(s) full responsibility for the content of the published article.

## REFERENCES

1. Qin, S. et al. mRNA-based therapeutics: powerful and versatile tools to combat diseases. Signal Transduction and Targeted Therapy 7, 166 (2022).

2. Sahin, U., Karikó, K. & Türeci, Ö. mRNA-based therapeutics--developing a new class of drugs. Nat. Rev. Drug Discov. 13, 759–780 (2014).

3. Rohner, E., Yang, R., Foo, K. S., Goedel, A. & Chien, K. R. Unlocking the promise of mRNA therapeutics. Nat. Biotechnol. 40, 1586–1600 (2022).

4. Roberts, T. C., Langer, R. & Wood, M. J. A. Advances in oligonucleotide drug delivery. Nat. Rev. Drug Discov. 19, 673–694 (2020).

5. Metkar, M., Pepin, C. S. & Moore, M. J. Tailor made: the art of therapeutic mRNA design. Nat. Rev. Drug Discov. 23, 67–83 (2024).

6. Baden, L. R. et al. Efficacy and safety of the mRNA-1273 SARS-CoV-2 vaccine. N. Engl. J. Med. 384, 403–416 (2021).

7. Walsh, E. E. et al. Safety and immunogenicity of two RNA-based Covid-19 vaccine candidates. N. Engl. J. Med. 383, 2439–2450 (2020).

8. Yaremenko, A. V., Khan, M. M., Zhen, X., Tang, Y. & Tao, W. Clinical advances of mRNA vaccines for cancer immunotherapy. Med (N. Y.) 6, 100562 (2025).

9. Mauger, D. M. et al. mRNA structure regulates protein expression through changes in functional half-life. Proc. Natl. Acad. Sci. U. S. A. 116, 24075–24083 (2019).

10. Leppek, K. et al. Combinatorial optimization of mRNA structure, stability, and translation for RNA-based therapeutics. Nat. Commun. 13, 1536 (2022).

11. Chen, R. et al. Engineering circular RNA for enhanced protein production. Nat. Biotechnol. 41, 262–272 (2023).

12. Chen, C.-K. et al. Structured elements drive extensive circular RNA translation. Mol. Cell 81, 4300–4318.e13 (2021).

13. Jin, L., Zhou, Y., Zhang, S. & Chen, S.-J. mRNA vaccine sequence and structure design and optimization: Advances and challenges. J. Biol. Chem. 301, 108015 (2025).

14. Castillo-Hair, S. et al. Optimizing 5’UTRs for mRNA-delivered gene editing using deep learning. Nat. Commun. 15, 5284 (2024).

15. Yoon, S. et al. Designing 5’ UTR sequences improves the capacity of mRNA therapeutics in preclinical models of aging and obesity. Mol. Ther. (2026) doi:10.1016/j.ymthe.2025.12.060.

16. Sample, P. J. et al. Human 5’ UTR design and variant effect prediction from a massively parallel translation assay. Nat. Biotechnol. 37, 803–809 (2019).

17. Li, X., Yang, L. & Chen, L.-L. The biogenesis, functions, and challenges of circular RNAs. Mol. Cell 71, 428–442 (2018).

18. Chen, L.-L. The biogenesis and emerging roles of circular RNAs. Nat. Rev. Mol. Cell Biol. 17, 205–211 (2016).

19. Liu, C.-X. & Chen, L.-L. Circular RNAs: Characterization, cellular roles, and applications. Cell 185, 2016–2034 (2022).

20. Obi, P. & Chen, Y. G. The design and synthesis of circular RNAs. Methods 196, 85–103 (2021).

21. Barrett, S. P. & Salzman, J. Circular RNAs: analysis, expression and potential functions. Development 143, 1838–1847 (2016).

22. Kirio, K. et al. Circular RNAs exhibit exceptional stability in the aging brain and serve as reliable age and experience indicators. Cell Rep. 44, 115485 (2025).

23. Qu, L. et al. Circular RNA vaccines against SARS-CoV-2 and emerging variants. Cell 185, 1728–1744.e16 (2022).

24. Suo, J. et al. Circular RNA-based protein replacement therapy mitigates osteoarthritis in male mice. Nat. Commun. 16, 8480 (2025).

25. Wesselhoeft, R. A. et al. RNA Circularization Diminishes Immunogenicity and Can Extend Translation Duration In Vivo. Mol. Cell 74, 508–520.e4 (2019).

26. Wen, S.-Y., Qadir, J. & Yang, B. B. Circular RNA translation: novel protein isoforms and clinical significance. Trends Mol. Med. 28, 405–420 (2022).

27. Nielsen, A. F. et al. Best practice standards for circular RNA research. Nat. Methods 19, 1208–1220 (2022).

28. Pamudurti, N. R. et al. Translation of CircRNAs. Mol. Cell 66, 9–21 (2017).

29. Chen, Y. G. et al. Sensing self and foreign circular RNAs by intron identity. Mol. Cell 67, 228–238.e5 (2017).

30. Wesselhoeft, R. A., Kowalski, P. S. & Anderson, D. G. Engineering circular RNA for potent and stable translation in eukaryotic cells. Nat. Commun. 9, 2629 (2018).

31. Chen, H. et al. Chemical and topological design of multicapped mRNA and capped circular RNA to augment translation. Nat. Biotechnol. 43, 1128–1143 (2025).

32. Jackson, R. J., Hellen, C. U. T. & Pestova, T. V. The mechanism of eukaryotic translation initiation and principles of its regulation. Nat. Rev. Mol. Cell Biol. 11, 113–127 (2010).

33. Yang, Y. et al. Extensive translation of circular RNAs driven by N6-methyladenosine. Cell Res. 27, 626–641 (2017).

34. Jia, L. et al. Decoding mRNA translatability and stability from the 5’ UTR. Nat. Struct. Mol. Biol. 27, 814–821 (2020).

35. Wu, Q. & Bazzini, A. A. Translation and mRNA Stability Control. Annu. Rev. Biochem. 92, 227–245 (2023).

36. Johnson, A. G., Grosely, R., Petrov, A. N. & Puglisi, J. D. Dynamics of IRES-mediated translation. Philos. Trans. R. Soc. Lond. B Biol. Sci. 372, (2017).

37. Thompson, S. R. Tricks an IRES uses to enslave ribosomes. Trends Microbiol. 20, 558–566 (2012).

38. Weingarten-Gabbay, S. et al. Systematic discovery of cap-independent translation sequences in human and viral genomes. Science 351, 1–24 (2016).

39. Byeon, G. W. et al. Functional and structural basis of extreme conservation in vertebrate 5’ untranslated regions. Nat. Genet. 53, 729–741 (2021).

40. Leppek, K., Das, R. & Barna, M. Functional 5′ UTR mRNA structures in eukaryotic translation regulation and how to find them. Nature Reviews Molecular Cell Biology Preprint at 10.1038/nrm.2017.103 (2018).

41. Martinez-Salas, E., Francisco-Velilla, R., Fernandez-Chamorro, J. & Embarek, A. M. Insights into Structural and Mechanistic Features of Viral IRES Elements. Front. Microbiol. 8, 2629 (2017).

42. Brito Querido, J., Díaz-López, I. & Ramakrishnan, V. The molecular basis of translation initiation and its regulation in eukaryotes. Nat. Rev. Mol. Cell Biol. (2023) doi:10.1038/s41580-023-00624-9.

43. Komar, A. A. & Hatzoglou, M. Cellular IRES-mediated translation: the war of ITAFs in pathophysiological states. Cell Cycle 10, 229–240 (2011).

44. Leppek, K. et al. Gene- and species-specific Hox mRNA translation by ribosome expansion segments. Mol. Cell Nov 4;S1097–2765(20)30730–9 (2020) doi:10.1016/j.molcel.2020.10.023.

45. Leppek, K., Byeon, G. W., Fujii, K. & Barna, M. VELCRO-IP RNA-seq reveals ribosome expansion segment function in translation genome-wide. Cell Rep. 34, 108629 (2021).

46. Sherlock, M. E., Langeberg, C. J., Segar, K. E. & Kieft, J. S. A conserved class of viral RNA structures regulate translation reinitiation through dynamic ribosome interactions. bioRxiv (2023) doi:10.1101/2023.09.29.560040.

47. Jackson, R. J. The current status of vertebrate cellular mRNA IRESs. Cold Spring Harb. Perspect. Biol. 5, a011569, 1–20 (2013).

48. Holcik, M. & Sonenberg, N. Translational control in stress and apoptosis. Nat. Rev. Mol. Cell Biol. 6, 318–327 (2005).

49. Spriggs, K. A., Bushell, M. & Willis, A. E. Translational regulation of gene expression during conditions of cell stress. Mol. Cell 40, 228–237 (2010).

50. Koch, A., Aguilera, L., Morisaki, T., Munsky, B. & Stasevich, T. J. Quantifying the dynamics of IRES and cap translation with single-molecule resolution in live cells. Nat. Struct. Mol. Biol. 27, 1095–1104 (2020).

51. Cornelis, S. et al. Identification and characterization of a novel cell cycle-regulated internal ribosome entry site. Mol. Cell 5, 597–605 (2000).

52. Xue, S. et al. RNA regulons in Hox 5’ UTRs confer ribosome specificity to gene regulation. Nature 517, 33–38 (2015).

53. Oh, S.-K., Scott, M. P. & Sarnow, P. Homeotic gene Antennapedia mRNA confer translational initiation by internal ribosome binding. Genes Dev. 6, 1643–1653 (1992).

54. Ye, X., Fong, P., Iizuka, N., Choate, D. & Cavener, D. R. ultrabithorax and antennapedia 5′ untranslated regions promote developmentally regulated internal translation initiation. Mol. Cell. Biol. 17, 1714–1721 (1997).

55. Fan, X., Yang, Y., Chen, C. & Wang, Z. Pervasive translation of circular RNAs driven by short IRES-like elements. Nat. Commun. 13, 3751 (2022).

56. Chen, C. Y. & Sarnow, P. Initiation of protein synthesis by the eukaryotic translational apparatus on circular RNAs. Science 268, 415–417 (1995).

57. Hansen, T. B. et al. Natural RNA circles function as efficient microRNA sponges. Nature 495, 384–388 (2013).

58. Jeck, W. R. & Sharpless, N. E. Detecting and characterizing circular RNAs. Nat. Biotechnol. 32, 453–461 (2014).

59. Salzman, J., Gawad, C., Wang, P. L., Lacayo, N. & Brown, P. O. Circular RNAs are the predominant transcript isoform from hundreds of human genes in diverse cell types. PLoS One 7, e30733 (2012).

60. Salzman, J., Chen, R. E., Olsen, M. N., Wang, P. L. & Brown, P. O. Cell-type specific features of circular RNA expression. PLoS Genet. 9, e1003777 (2013).

61. Panda, A. C. et al. High-purity circular RNA isolation method (RPAD) reveals vast collection of intronic circRNAs. Nucleic Acids Res. 45, e116 (2017).

62. Zhang, Y. et al. The Biogenesis of Nascent Circular RNAs. Cell Rep. 15, 611–624 (2016).

63. Liu, C.-X., Yang, L. & Chen, L.-L. Dynamic conformation: Marching toward circular RNA function and application. Mol. Cell 84, 3596–3609 (2024).

64. Chen, L.-L. The expanding regulatory mechanisms and cellular functions of circular RNAs. Nat. Rev. Mol. Cell Biol. 21, 475–490 (2020).

65. Legnini, I. et al. Circ-ZNF609 Is a Circular RNA that Can Be Translated and Functions in Myogenesis. Mol. Cell 66, 22–37 (2017).

66. Liang, W.-C. et al. Translation of the circular RNA circβ-catenin promotes liver cancer cell growth through activation of the Wnt pathway. Genome Biol. 20, 84 (2019).

67. Yang, Y. et al. Novel Role of FBXW7 Circular RNA in Repressing Glioma Tumorigenesis. J. Natl. Cancer Inst. 110, 304–315 (2018).

68. Zhang, M. et al. A novel protein encoded by the circular form of the SHPRH gene suppresses glioma tumorigenesis. Oncogene 37, 1805–1814 (2018).

69. Chen, Y. G. et al. N6-methyladenosine modification controls circular RNA immunity. Mol. Cell 76, 96–109.e9 (2019).

70. Liu, C.-X. et al. Structure and degradation of circular RNAs regulate PKR activation in innate immunity. Cell 177, 865–880.e21 (2019).

71. Tai, J. & Chen, Y. G. Differences in the immunogenicity of engineered circular RNAs. J. Mol. Cell Biol. 15, (2023).

72. Liu, C.-X. et al. RNA circles with minimized immunogenicity as potent PKR inhibitors. Mol. Cell 82, 420–434.e6 (2022).

73. Hornung, V. et al. 5’-Triphosphate RNA is the ligand for RIG-I. Science 314, 994–997 (2006).

74. Kato, H. et al. Length-dependent recognition of double-stranded ribonucleic acids by retinoic acid-inducible gene-I and melanoma differentiation-associated gene 5. J. Exp. Med. 205, 1601–1610 (2008).

75. Loo, Y.-M. & Gale, M., Jr. Immune signaling by RIG-I-like receptors. Immunity 34, 680–692 (2011).

76. Liang, D. & Wilusz, J. E. Short intronic repeat sequences facilitate circular RNA production. Genes Dev. 28, 2233–2247 (2014).

77. Ford, E. & Ares, M., Jr. Synthesis of circular RNA in bacteria and yeast using RNA cyclase ribozymes derived from a group I intron of phage T4. Proc. Natl. Acad. Sci. U. S. A. 91, 3117–3121 (1994).

78. Perriman, R. & Ares, M., Jr. Circular mRNA can direct translation of extremely long repeating-sequence proteins in vivo. RNA 4, 1047–1054 (1998).

79. Petkovic, S. & Müller, S. RNA circularization strategies in vivo and in vitro. Nucleic Acids Res. 43, 2454–2465 (2015).

80. Chen, C., Yang, Y. & Wang, Z. Study of circular RNA translation using reporter systems in living cells. Methods 196, 113–120 (2021).

81. Koch, P. et al. A versatile toolbox for determining IRES activity in cells and embryonic tissues. EMBO J. (2025) doi:10.1038/s44318-025-00404-5.

82. Arendrup, F. S. W., Andersen, K. L. & Lund, A. H. A tripartite cell-free translation system to study mammalian translation. Nat. Protoc. 20, 2803–2844 (2025).

83. Roberts, B. E. & Paterson, B. M. Efficient translation of tobacco mosaic virus RNA and rabbit globin 9S RNA in a cell-free system from commercial wheat germ. Proc. Natl. Acad. Sci. U. S. A. 70, 2330–2334 (1973).

84. Carlson, E. D., Gan, R., Hodgman, C. E. & Jewett, M. C. Cell-free protein synthesis: applications come of age. Biotechnol. Adv. 30, 1185–1194 (2012).

85. Soto Rifo, R., Ricci, E. P., Décimo, D., Moncorgé, O. & Ohlmann, T. Back to basics: the untreated rabbit reticulocyte lysate as a competitive system to recapitulate cap/poly(A) synergy and the selective advantage of IRES-driven translation. Nucleic Acids Res. 35, e121 (2007).

86. Bergamini, G. & Gebauer, F. Poly(A)-dependent cell-free translation systems from animal cells. in Cell-Free Translation Systems 79–88 (Springer Berlin Heidelberg, Berlin, Heidelberg, 2002). doi:10.1007/978-3-642-59379-6_7.

87. Kondrashov, N. et al. Ribosome-mediated specificity in Hox mRNA translation and vertebrate tissue patterning. Cell 145, 383–397 (2011).

88. Akirtava, C., May, G. E. & McManus, C. J. False-positive IRESes from Hoxa9 and other genes resulting from errors in mammalian 5’ UTR annotations. Proc. Natl. Acad. Sci. U. S. A. 119, e2122170119 (2022).

89. Stoneley, M., Paulin, F. E. M., Quesne, J. P. C. L., Chappell, S. A. & Willis, A. E. C-Myc 5′ untranslated region contains an internal ribosome entry segment. Oncogene 16, 423–428 (1998).

90. Nanbru, C. et al. Alternative translation of the proto-oncogene c-mycby an internal ribosome entry site. J. Biol. Chem. 272, 32061–32066 (1997).

91. Choi, J.-H. et al. IRES-mediated translation of cofilin regulates axonal growth cone extension and turning. EMBO J. 37, (2018).

92. Fitzgerald, K. D. & Semler, B. L. Bridging IRES elements in mRNAs to the eukaryotic translation apparatus. Biochim. Biophys. Acta 1789, 518–528 (2009).

93. Thompson, S. R. So you want to know if your message has an IRES? Wiley Interdisciplinary Reviews: RNA vol. 3 697–705 Preprint at 10.1002/wrna.1129 (2012).

94. Lindley, S. R. et al. Ribozyme-activated mRNA trans-ligation enables large gene delivery to treat muscular dystrophies. Science 386, 762–767 (2024).

95. Unti, M. J. & Jaffrey, S. R. Highly efficient cellular expression of circular mRNA enables prolonged protein expression. Cell Chem. Biol. 31, 163–176.e5 (2024).

96. Jiang, Y., Chen, X. & Zhang, W. Overexpression-based detection of translatable circular RNAs is vulnerable to coexistent linear RNA byproducts. Biochem. Biophys. Res. Commun. 558, 189–195 (2021).

97. Ho-Xuan, H. et al. Comprehensive analysis of translation from overexpressed circular RNAs reveals pervasive translation from linear transcripts. Nucleic Acids Res. 48, 10368–10382 (2020).

98. Litke, J. L. & Jaffrey, S. R. Highly efficient expression of circular RNA aptamers in cells using autocatalytic transcripts. Nat. Biotechnol. 37, 667–675 (2019).

99. Roth, A. et al. A widespread self-cleaving ribozyme class is revealed by bioinformatics. Nat. Chem. Biol. 10, 56–60 (2014).

100. Saito, T., Owen, D. M., Jiang, F., Marcotrigiano, J. & Gale, M., Jr. Innate immunity induced by composition-dependent RIG-I recognition of hepatitis C virus RNA. Nature 454, 523–527 (2008).

101. Quade, N., Boehringer, D., Leibundgut, M., van den Heuvel, J. & Ban, N. Cryo-EM structure of Hepatitis C virus IRES bound to the human ribosome at 3.9-Å resolution. Nat. Commun. 6, 7646 (2015).

102. Yamamoto, H. et al. Molecular architecture of the ribosome-bound Hepatitis C Virus internal ribosomal entry site RNA. EMBO J. 34, 3042–3058 (2015).

103. Jaafar, Z. A., Oguro, A., Nakamura, Y. & Kieft, J. S. Translation initiation by the hepatitis C virus IRES requires eIF1A and ribosomal complex remodeling. Elife 5, (2016).

104. Kieft, J. S., Zhou, K., Jubin, R. & Doudna, J. A. Mechanism of ribosome recruitment by hepatitis C IRES RNA. RNA 7, 194–206 (2001).

105. Panganiban, G. & Rubenstein, J. L. R. Developmental functions of the Distal-less/Dlx homeobox genes. Development 129, 4371–4386 (2002).

106. Depew, M. J., Simpson, C. A., Morasso, M. & Rubenstein, J. L. R. Reassessing the Dlx code: the genetic regulation of branchial arch skeletal pattern and development. J. Anat. 207, 501–561 (2005).

107. Meyer, K. D. et al. 5’ UTR m(6)A promotes cap-independent translation. Cell 163, 999–1010 (2015).

108. Huang, Y. et al. IRES-cargo interplay structurally modulates circular RNA translation. Cell Res. (2026) doi:10.1038/s41422-026-01233-9.

109. Liao, K.-C. et al. Characterization of group I introns in generating circular RNAs as vaccines. Nucleic Acids Res. 53, gkaf089 (2025).

110. Osuna, B. A., Howard, C. J., Kc, S., Frost, A. & Weinberg, D. E. In vitro analysis of RQC activities provides insights into the mechanism and function of CAT tailing. Elife 6, (2017).

