## Appendix for "Synthetic circRNAs employ IRES activity for translation in cells and in cell-free translation systems"

### Table of Contents

**This document includes:** page

Appendix Figure S1 to S5 ........ 2-11
Appendix Table S1 ........ 12-13
Appendix Table S2 ........ 14-15

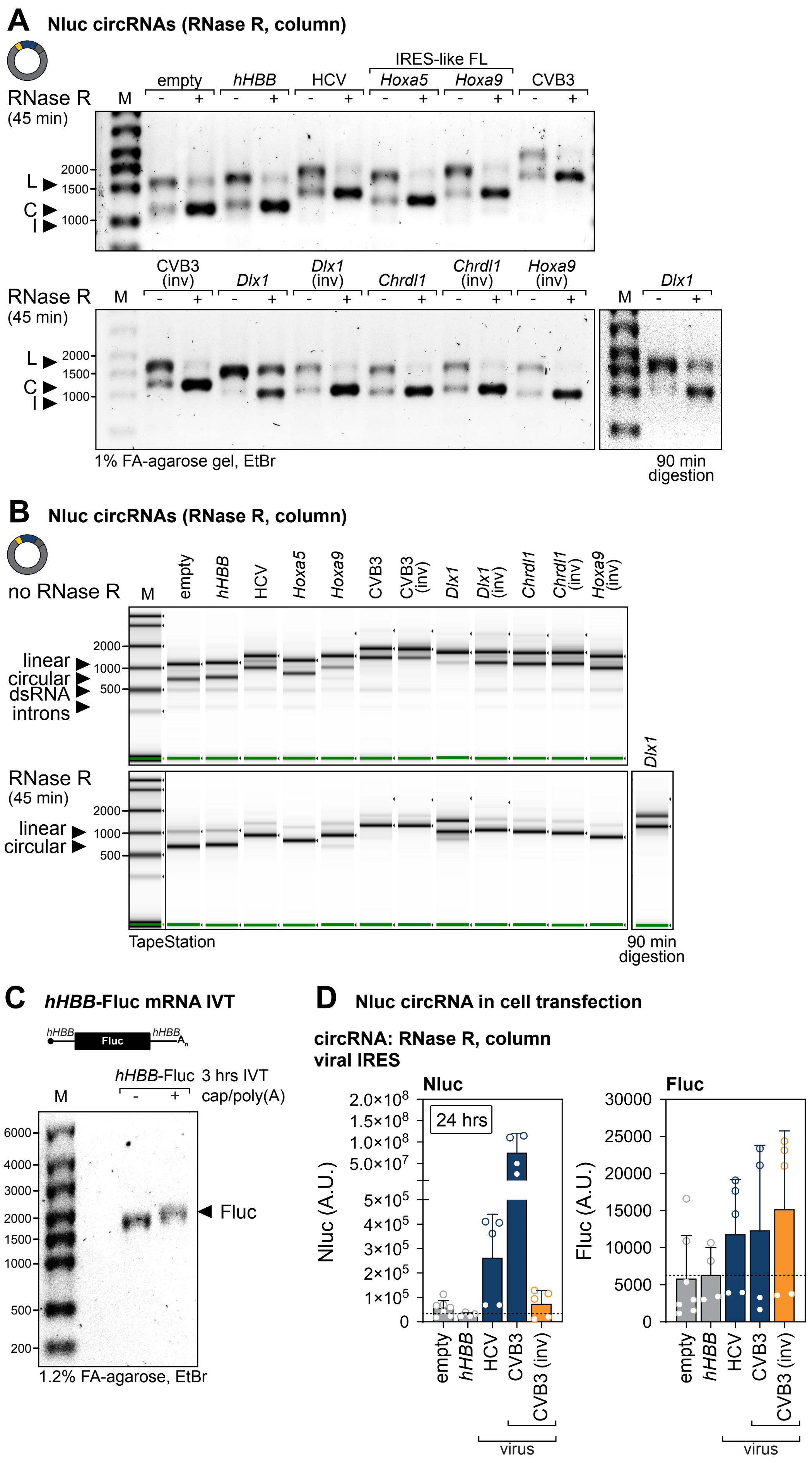
A
Nluc circRNAs (RNase R, column)
RNase R
(45 min)
M
empty
hHBB
HCV
IRES-like FL
Hoxa5
Hoxa9
CVB3
- +
L
C
I
2000
1500
1000
CVB3 (inv)
Dlx1
Dlx1 (inv)
Chrdl1
Chrdl1 (inv)
Hoxa9 (inv)
1% FA-agarose gel, EtBr
90 min digestion
B
Nluc circRNAs (RNase R, column)
no RNase R
linear
circular
dsRNA
introns
500
TapeStation
C
hHBB-Fluc mRNA IVT
hHBB
Fluc
hHBB-Fluc 3 hrs IVT
cap/poly(A)
6000
4000
3000
200
1.2% FA-agarose, EtBr
D
Nluc circRNA in cell transfection
circRNA: RNase R, column
viral IRES
Nluc
24 hrs
Nluc (A.U.)
2.0×10^8
1.5×10^8
1.0×10^8
5.0×10^7
5×10^5
4×10^5
3×10^5
2×10^5
1×10^5
0
CVB3 (inv)
virus
Fluc (A.U.)
30000
25000
20000
15000
10000
5000

**Appendix Figure S1. circRNA reporter *in vitro* transcription and RNase R purification.**

(**A**) Quality control of the generated 3xHA-Nluc encoding circRNAs using 1% FA-agarose gel. Linear side products (upper band) are mainly degraded by RNase R digestion. Only the circular RNase R-resistant band remains (lower band). Only *Dlx1* shows less efficient circularization leading to a higher concentration of side products. Therefore, the RNase R incubation time was increased to 90 min. 500 ng total RNA was loaded per lane. RNA species: linear (L), circular (C), introns (I).

(**B**) Quality control, using the High Sensitivity RNA ScreenTape, of the generated Nluc circRNAs before purification (upper panel) and after RNase R digestion and column purification (lower panel). Remaining contaminants can be observed as light grey bands. Full gel represented as in **Fig. 1D** and **3B**.

(**C**) IVT RNA and gel extracted, RNase R treated, and column purified circRNA was analyzed using the Agilent RNA high sensitivity screen tape. The corresponding electrogram data of the unpurified IVT RNA (grey) as well as the RNase R and gel-purified circRNA (colour-coded according to the tested IRES insert) are overlaid. Intron sequences, linear and circular RNA species are indicated according to size.

(**D**) IRES activity was evaluated 24 hpt using Nluc circRNAs and *hHBB*-Fluc linear capped control RNA, corresponding to Nluc/Fluc ratios normalized to empty control in **Fig. 1F**. Bar graphs present the raw Nluc and raw Fluc data; n= 3-5.

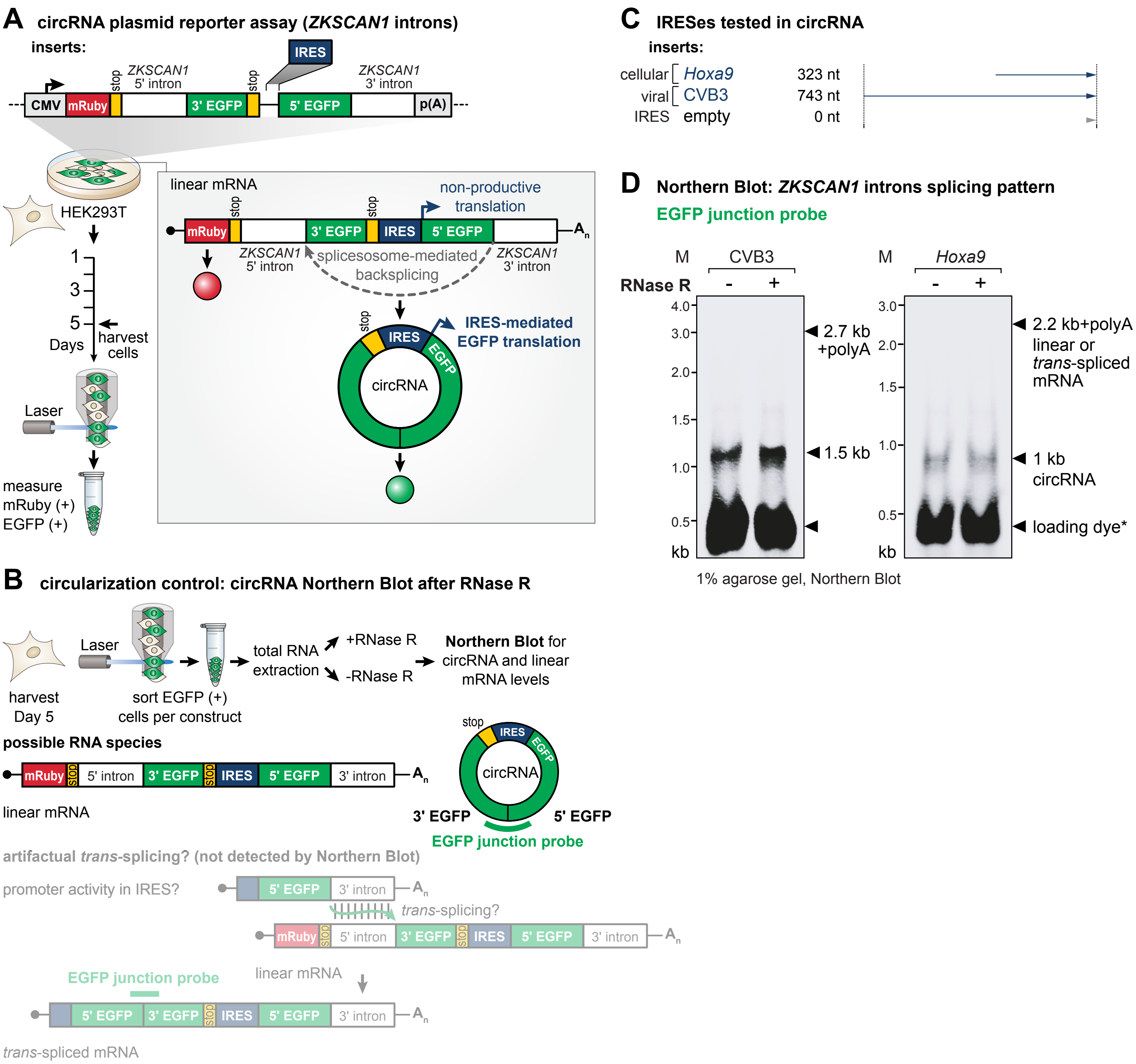
A circRNA plasmid reporter assay (ZKSCAN1 introns)
inserts:
IRES
stop
ZKSCAN1 5' intron
ZKSCAN1 3' intron
CMV
mRuby
3' EGFP
5' EGFP
p(A)
HEK293T
1
3
5
Days
harvest cells
Laser
measure mRuby (+) EGFP (+)
linear mRNA
non-productive translation
A
n
splicesosome-mediated backsplicing
IRES-mediated EGFP translation
circRNA
EGFP
B circularization control: circRNA Northern Blot after RNase R
Laser
harvest Day 5
sort EGFP (+) cells per construct
total RNA extraction
+RNase R
-RNase R
Northern Blot for circRNA and linear mRNA levels
possible RNA species
5' intron
3' intron
linear mRNA
3' EGFP
5' EGFP
EGFP junction probe
artifactual trans-splicing? (not detected by Northern Blot)
promoter activity in IRES?
trans-splicing?
trans-spliced mRNA
C IRESes tested in circRNA
inserts:
cellular Hoxa9 323 nt
viral CVB3 743 nt
IRES empty 0 nt
D Northern Blot: ZKSCAN1 introns splicing pattern
EGFP junction probe
M
CVB3
Hoxa9
RNase R
-
+
4.0
3.0
2.0
1.5
1.0
0.5
kb
2.7 kb +polyA
1.5 kb
2.2 kb+polyA linear or trans-spliced mRNA
1 kb circRNA
loading dye*
1% agarose gel, Northern Blot

**Appendix Figure S2. Northern blot analysis of the ZKSCAN1-based back-spliced plasmid reporter system for circRNA generation in cells.**

(**A**) Experimental outline of the circRNA reporter assay based on the mRuby-ZKSCAN1-splitEGFP plasmid for testing of the IRES activity of different IRES sequences. Following plasmid transfection, the expression of the reporter system under CMV promoter control leads to the linear pre-mRNA which is circularized through spliceosome-mediated back-splicing (grey box). Cells were harvested at 5 dpt, and their mRuby signal (transfection control) and EGFP signal (readout for IRES activity) is detected. Schematic partially adapted from (Chen *et al*, 2021).

(**B**) Schematic of the circularization control with the mRuby-ZKSCAN1-splitEGFP plasmid system by Northern blot. For this, EGFP+ cells are sorted at 5 dpt and total RNA is treated with and without RNase R to distinguish linear mRNA and circRNA levels. The EGFP junction probe used for Northern blot is indicated along with possible RNA species expressed and which of those the probe would detect. The EGFP junction probe should detect mature circRNA, but not pre-mRNA, and is also sensitive for potential *trans*-spliced mRNA species.

(**C**) We test a representative viral (CVB3) and cellular (*Hoxa9*) IRES in the mRuby-ZKSCAN1-splitEGFP plasmid system.

(**D**) Northern blot analysis of the mRuby-ZKSCAN1-splitEGFP circRNA reporter transfected and mRuby+/EGFP+ sorted cells using probes against the EGFP junction region on the reporter transcript with or without RNase R treatment. The EGFP circRNA reporter construct does not generate an EGFP signal corresponding to linear pre-mRNAs or *trans*-spliced mRNAs, but exclusively for circRNAs.

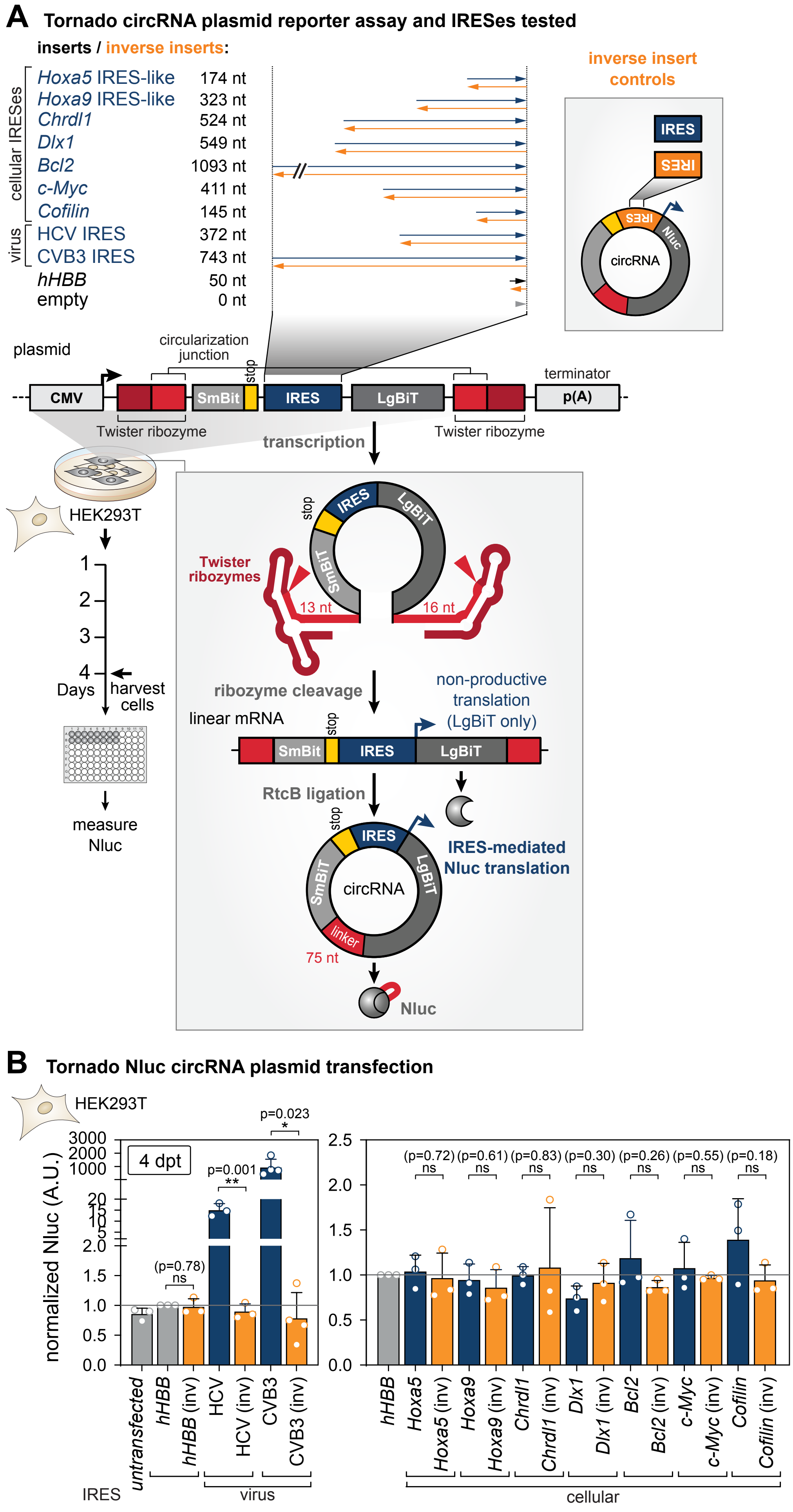
A Tornado circRNA plasmid reporter assay and IRESes tested
inserts / inverse inserts:
cellular IRESes
Hoxa5 IRES-like 174 nt
Hoxa9 IRES-like 323 nt
Chrdl1 524 nt
Dlx1 549 nt
Bcl2 1093 nt
c-Myc 411 nt
Cofilin 145 nt
virus
HCV IRES 372 nt
CVB3 IRES 743 nt
hHBB 50 nt
empty 0 nt
inverse insert controls
IRES
circRNA
Nluc
plasmid
circularization junction
stop
CMV
SmBit
IRES
LgBiT
terminator
p(A)
Twister ribozyme
Twister ribozyme
transcription
HEK293T
1
2
3
4
Days
harvest cells
measure Nluc
Twister ribozymes
13 nt
16 nt
ribozyme cleavage
non-productive translation (LgBiT only)
linear mRNA
RtcB ligation
IRES-mediated Nluc translation
linker
75 nt
Nluc
B Tornado Nluc circRNA plasmid transfection
HEK293T
4 dpt
normalized Nluc (A.U.)
(p=0.78) ns
p=0.001 **
p=0.023 *
(p=0.72) ns
(p=0.61) ns
(p=0.83) ns
(p=0.30) ns
(p=0.26) ns
(p=0.55) ns
(p=0.18) ns
untransfected
hHBB
hHBB (inv)
HCV
HCV (inv)
CVB3
CVB3 (inv)
Hoxa5
Hoxa5 (inv)
Hoxa9
Hoxa9 (inv)
Chrdl1
Chrdl1 (inv)
Dlx1
Dlx1 (inv)
Bcl2
Bcl2 (inv)
c-Myc
c-Myc (inv)
Cofilin
Cofilin (inv)
IRES
virus
cellular

**Appendix Figure S3. IRES-dependent circRNA translation of the plasmid-based self-splicing ribozyme-based Tornado translation system for circRNA generation in cells.**

(**A**) Experimental outline of the plasmid-based Tornado translation system that generates Nluc circRNA reporters by Twister ribozyme cleavage and RtcB ligation. We test the IRES activity of seven different cellular IRES and two viral IRESes, and their respective inverse controls. Following plasmid transfection, the expression of the reporter system under CMV promoter control leads to the linear pre-mRNA which is circularized through ribozyme activity (grey box). Cells were harvested at 4 dpt, and their Nluc signal (readout for IRES activity) is detected. Schematic partially adapted from (Unti *et al*, 2024).

(**B**) IRES activity of the tested sequences was evaluated 4 dpt using 2 µg Tornado Nluc circRNA plasmids transfected with 2 µL Lipofectamine 3000 into HEK203T cells in a 12-well plate. Tested IRES sequences are depicted in dark blue, inverse controls in orange, negative controls in grey. Activities of IRESes of viral and cellular origin are depicted separately, with the same *hHBB* data included in both viral and cellular panels. Nluc data (A.U.) are normalized to the *hHBB* control, set to 1. Bar graphs are indicating mean values ± SD, n = 3-4; ns, not significant.

**circRNA IVT analysis after urea-PAGE gel-extraction, RNase R**

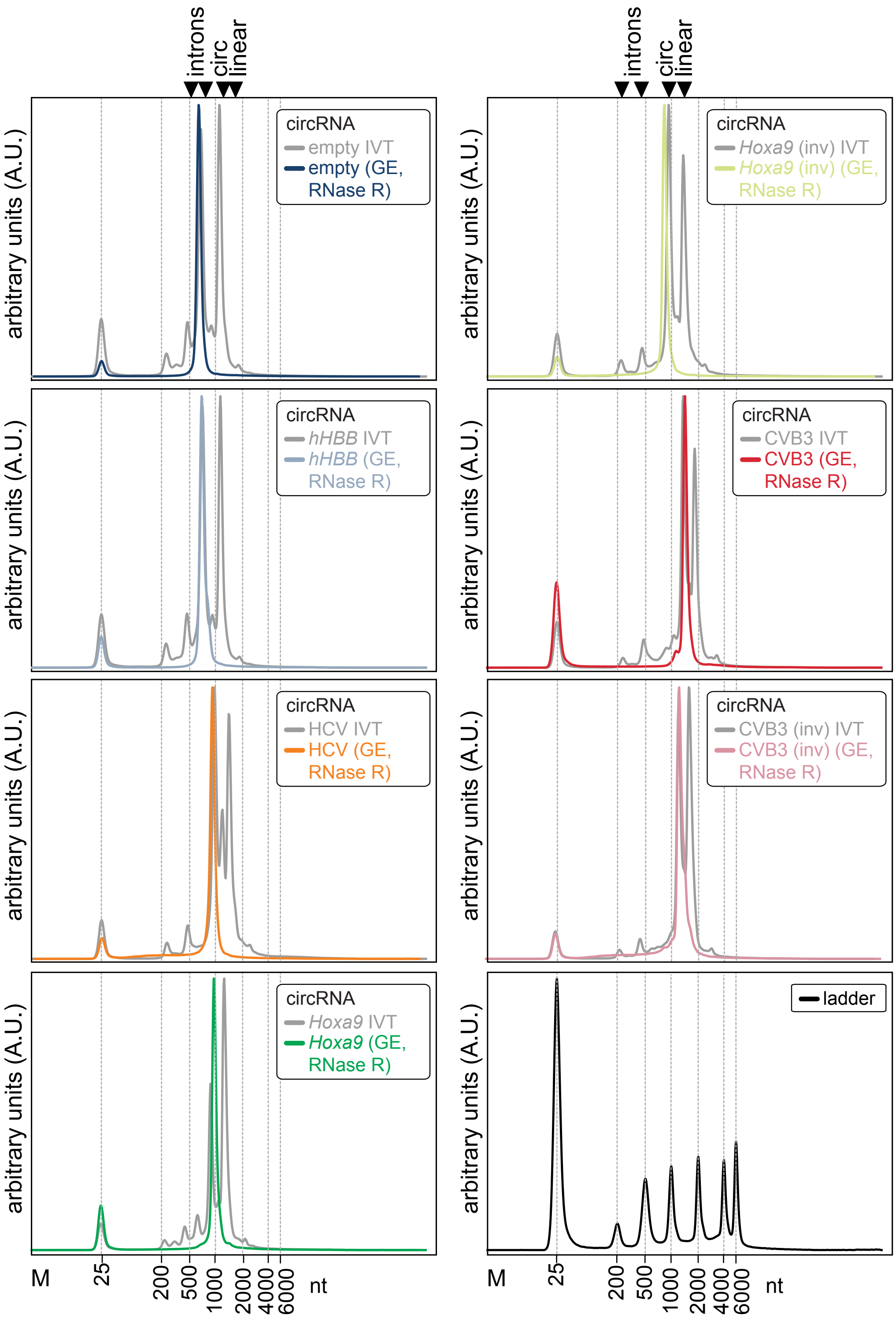

**Appendix Figure S4. Qualitative analysis of hHBB-Fluc control mRNA and of urea-PAGE gel extracted circRNAs after extraction, RNase R, and column purification.**

(**A**) Quality control of the mRNA *hHBB*-Fluc used as a co-transfection control. mRNA was analyzed on a 1.2% formaldehyde (FA) gel stained with ethidium bromide (EtBr) after 3 hrs of *in vitro* transcription (IVT) and enzymatic capping and polyadenylation. The RiboRuler High Range RNA ladder (Thermo Fisher) is loaded for reference; molecular weight given in base-pairs. This result has been repeated independently >3 times with similar results.

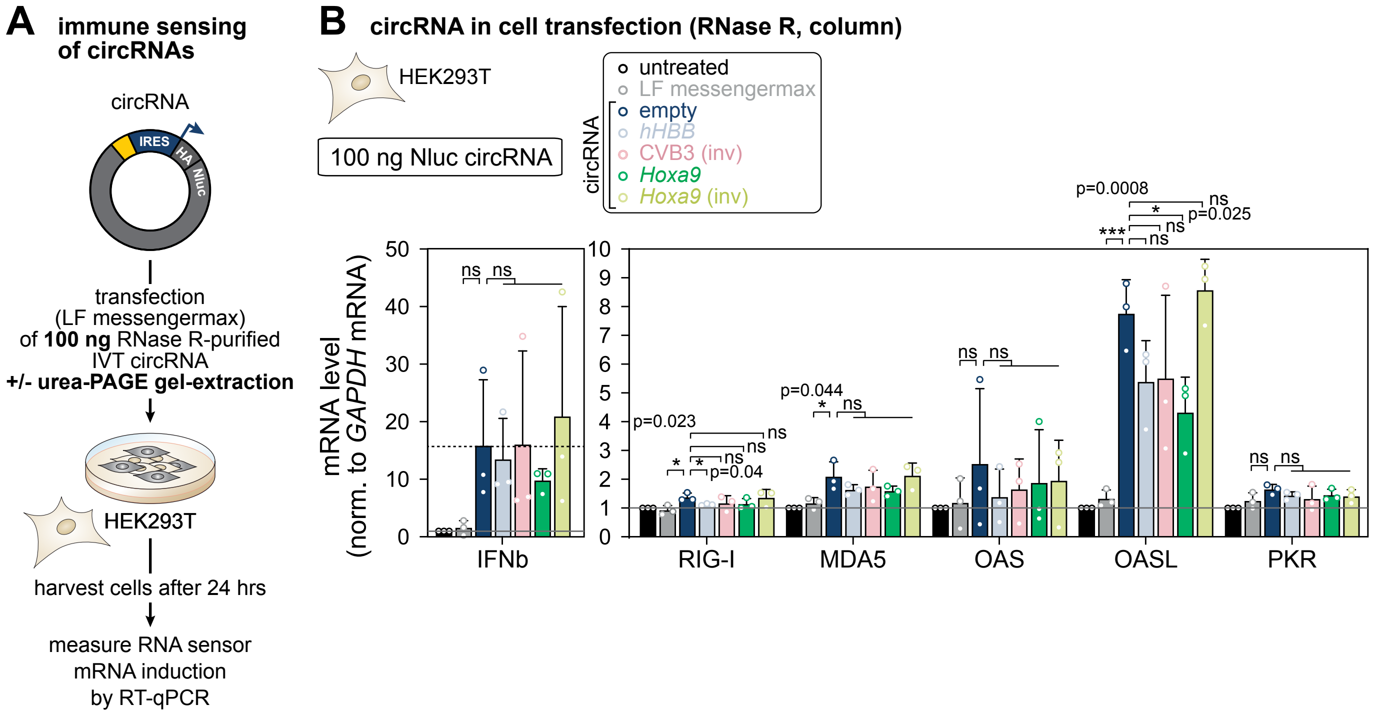
A
immune sensing of circRNAs
circRNA
IRES
HA
Nluc
transfection (LF messengermax) of 100 ng RNase R-purified IVT circRNA
+/- urea-PAGE gel-extraction
HEK293T
harvest cells after 24 hrs
measure RNA sensor mRNA induction by RT-qPCR
B
circRNA in cell transfection (RNase R, column)
HEK293T
100 ng Nluc circRNA
untreated
LF messengermax
circRNA
empty
hHBB
CVB3 (inv)
Hoxa9
Hoxa9 (inv)
mRNA level (norm. to GAPDH mRNA)
IFNb
RIG-I
MDA5
OAS
OASL
PKR
p=0.023
p=0.04
p=0.044
p=0.0008
p=0.025

### Appendix Figure S5. circRNA sensing and evaluation of IRES-specific immunogenicity of circRNAs.

(**A**) Experimental outline of the immune sensing experiment evaluating the immunogenicity of 100 ng transfected Nluc circRNAs containing different IRES sequences after purification. RNA was purified by RNase R only and column purification. HEK293T cells were harvested 24 hpt and RNA sensing was evaluated by RT-qPCR quantification of immune sensor RNA expression normalized to human GAPDH mRNA levels.

(**B**) Evaluation of purified circRNA immunogenicity dependent on encoded IRES sequences (indicated by color) 24 hpt. Transfected circRNA was RNase R digested and column purified. mRNA expression levels of different immune sensors were quantified (normalized to human GAPDH mRNA levels) after 100 ng circRNA transfection. Colors of the bar graphs according to the tested IRES sequences. mRNA expression levels are shown relative to untreated cells. Bar graphs are indicating mean values ± SD, n = 3. Data for untreated and *Hoxa9* is included as examples as well in **Fig 4F**.

### APPENDIX TABLES

#### Appendix Table S1: Plasmids used in this study.

All plasmids used for mammalian transient transfection are listed in the table.

| Appendix Table S1. List of plasmids | | |
|---|---|---|
| **Plasmid** | **Notes** | **Reference** |
| ***Mammalian cells*** | | |
| **Expression constructs** | | |
| mRuby3-ZK-spEGFP | pKL477, kindly provided by C. K. Chen | (Chen et al. 2021) |
| mRuby3-ZK-spEGFP-CVB3 | pPK008 | (Koch et al. 2025) |
| mRuby3-ZK-spEGFP-Hoxa9 | pLL0013 | (Koch et al. 2025) |
| pcDNA3.1-5'UTR-3xHA-Nluc | pKL401, kindly provided by C. Howard | (Osuna et al. 2017) |
| ***In vitro transcription*** | | |
| **circRNA EGFP constructs** | | |
| circEGFP-ScaI-v3 | pKL480 | (Chen et al. 2021) |
| mRuby3_circ_EGFP_HBB | pLL001 | This study |
| mRuby3_circ_EGFP_HCV | pLL002 | This study |
| **circRNA Nluc constructs** | | |
| mRuby_circ 3xHA_NanoLuc_empty | pPK001 | (Chen et al. 2021) |
| mRuby_circ 3xHA_NanoLuc_HBB | pPK002 | This study |
| mRuby_circ 3xHA_NanoLuc_HCV | pPK003 | This study |
| mRuby_circ 3xHA_NanoLuc_Hoxa5(IRES) | pPK004 | This study |
| mRuby_circ 3xHA_NanoLuc_Hoxa9(IRES) | pPK005 | This study |
| mRuby_circ 3xHA_NanoLuc_CVB3 | pPK042 | This study |
| mRuby_circ 3xHA_NanoLuc_CVB3inv | pPK043 | This study |
| mRuby_circ 3xHA_NanoLuc_Dlx1 | pPK044 | This study |
| mRuby_circ 3xHA_NanoLuc_Dlx1inv | pPK045 | This study |
| mRuby_circ 3xHA_NanoLuc_Chrdl1 | pPK046 | This study |
| mRuby_circ 3xHA_NanoLuc_Chrdl1inv | pPK047 | This study |
| mRuby_circ 3xHA_NanoLuc_HCVdIId_CCC | pPK076 | This study |
| mRuby_circ 3xHA_NanoLuc_HCV dII | pPK077 | This study |
| mRuby_circ 3xHA_NanoLuc_c-Myc | pPK078 | This study |
| mRuby_circ 3xHA_NanoLuc_Cofilin | pPK079 | |
| **Tornado circRNA constructs** | | |
| pcDNA3.1+-Tornado-split-nLuc | Addgene (ID 212611) | (Unti et al. 2024) |
| Tornado-split-nLuc_HBB | pPK055 | This study |
| Tornado-split-nLuc_HBBinv | pPK056 | This study |
| Tornado-split-nLuc_HCV | pPK057 | This study |
| Tornado-split-nLuc_HCVinv | pPK058 | This study |
| Tornado-split-nLuc_CVB3 | pPK059 | This study |
| Tornado-split-nLuc_CVB3inv | pPK060 | This study |

| | | |
|---|---|---|
| Tornado-split-nLuc_Chrdl1 | pPK061 | This study |
| Tornado-split-nLuc_Chrdl1inv | pPK062 | This study |
| Tornado-split-nLuc_Dlx1 | pPK063 | This study |
| Tornado-split-nLuc_Dlx1inv | pPK064 | This study |
| Tornado-split-nLuc_Bcl2 | pPK065 | This study |
| Tornado-split-nLuc_Bcl2inv | pPK066 | This study |
| Tornado-split-nLuc_c-Myc | pPK067 | This study |
| Tornado-split-nLuc_c-Mycinv | pPK068 | This study |
| Tornado-split-nLuc_Cofilin | pPK069 | This study |
| Tornado-split-nLuc_Cofilininv | pPK070 | This study |
| Tornado-split-nLuc_Hoxa9 | pPK071 | This study |
| Tornado-split-nLuc_Hoxa9inv | pPK072 | This study |
| Tornado-split-nLuc_Hoxa5 | pPK073 | This study |
| Tornado-split-nLuc_Hoxa5inv | pPK074 | This study |

### Appendix Table S2: DNA Oligonucleotides used in this study.

All DNA oligonucleotides used for cloning and RT-qPCR are listed in the table. F, forward primer; R, reverse primer.

**Appendix Table S2. DNA oligonucleotides**

| Name | Sequence | Description |
|---|---|---|
| | ***qPCR primer*** | |
| PK352 | TGTGGGCAATGTCATCAAAA | human *RIG-I* F |
| PK353 | GAAGCACTTGCTACCTCTTGC | human *RIG-I* R |
| PK354 | GGCACCATGGGAAGTGATT | human *MDA5* F |
| PK355 | ATTTGGTAAGGCCTGAGCTG | human *MDA5* R |
| PK356 | GCTCCTACCCTGTGTGTGTGT | human *OAS* F |
| PK357 | TGGTGAGAGTACTGAGGAAGA | human *OAS* R |
| PK358 | AGGGTACAGATGGGACATCG | human *OASL* F |
| PK359 | AAGGGTTCACGATGAGGTTG | human *OASL* R |
| PK360 | TCGCTGGTATCACTCGTCTG | human *PKR* F |
| PK361 | GATTCTGAAGACCGCCAGAG | human *PKR* R |
| PK362 | CTCTCCTGTTGTGCTTCTCC | human *IFNb* F |
| PK363 | GTCAAAGTTCATCCTGTCCTTG | human *IFNb* R |
| PK364 | GTCTCCTCTGACTTCAACAGCG | human *GAPDH* F |
| PK365 | ACCACCCTGTTGCTGTAGCCAA | human *GAPDH* R |
| | ***T7 promoter IVT primer set*** | |
| LL078 | AAAAAAAAAAAAAAAAAAAAAGGCCAGTGAATTGTAATACGACTCACTATAGGGgggaattctagaga aaatttcgtctg | circRNA-IVT-PolyA-T7Promoter-F |
| LL079 | TTTTTTTTTTTTTTTTTTTTTTTTTTTTTTTTTTTctgcaggtcgactctagagaaag | circRNA-IVT-PolyT-R |
| | ***Cloning of Nluc-circRNA reporter plasmids*** | |
| PK001 | catatgccagtgttatggatatcaaaATGGCCGTTTACCCATACG | 3HANano_F |
| PK002 | catatgccagtgttatggatatcaaaATGGCCGTTTACCCATACG | 3HANano_R |
| PK003 | GCATTCTGGCGTAAagtacatttgcttctgacacaactgt | HBB_F |
| PK004 | caagaaaacatctactgagAGTggtgtctgtttg | HBB_R |
| PK005 | GCATTCTGGCGTAAagtTTGGGGGCGACACT | HCV_F |
| PK006 | caagaaaacatctactgagAGTTGTTACGTTTGGTTTTTCTTTGAGGT | HCV_R |
| PK007 | GCATTCTGGCGTAAagtatcaggcaggatttacgactg | A5_F |
| PK008 | caagaaaacatctactgagAGTtgcttgatttgt | A5_R |
| PK009 | GCATTCTGGCGTAAagtttgatcttttaatcttcgttggcc | A9_F |
| PK010 | caagaaaacatctactgagAGTtgcagtagcccg | A9_R |
| PK268 | GCATTCTGGCGTAAagtTTAAAACAGCCTGTGGGTTGATCCC | CVB3_F |
| PK269 | caagaaaacatctactgagAGTGGTTTGCTGTATTCAACTTAACAATGAATTGTAATGT | CVB3_R |
| PK270 | CGAACGCATTCTGGCGTAAagtGGTTTGCTGTATTCAACTTAACAATGAATTGTAATGT | invCVB3_F |
| PK271 | acccaagaaaacatctactgagAGTTTAAAACAGCCTGTGGGTTGATCCC | invCVB3_R |
| PK272 | CGAACGCATTCTGGCGTAAagtCAGGCGTTGGGGGCG | Dlx1_F |
| PK273 | acccaagaaaacatctactgagAGTCTCTTCTCGCGGGGTCTG | Dlx1_R |
| PK274 | CGAACGCATTCTGGCGTAAagtCTCTTCTCGCGGGGTCTGG | invDlx1_F |
| PK275 | acccaagaaaacatctactgagAGTCAGGCGTTGGGGGCG | invDlx1_R |
| PK276 | CGAACGCATTCTGGCGTAAagtGTGTGGTGGGGGCGC | Chrdl1_F |
| PK277 | acccaagaaaacatctactgagAGTCTTCTACTTTTTTCTCCTTCGAGCTACTG | Chrdl1_R |
| PK278 | CGAACGCATTCTGGCGTAAagtCTTCTACTTTTTTCTCCTTCGAGCTACTGC | invChrdl1_F |

| | | |
|---|---|---|
| PK279 | acccaagaaaacatctactgagAGTGTGTGGTGGGGGCGC | invChrdl1_R |
| PK280 | CGAACGCATTCTGGCGTAAagttgcagtagcccgcgc | invA9_F |
| PK281 | acccaagaaaacatctactgagAGTttgatcttttaatcttcgttggccacaattaaaaca | invA9_R |
| PK438 | CGAACGCATTCTGGCGTAAagtACTAGAACTCGCTGTAGTAATTCCAGCG | cMyc_F |
| PK439 | acccaagaaaacatctactgagAGTTCGCGGGAGGCTGC | cMyc_R |
| PK440 | CGAACGCATTCTGGCGTAAagtGCCGGAAGGCCGCC | Cofilin_F |
| PK441 | acccaagaaaacatctactgagAGTGTTTCCGGAAACGAAAGGGAGAC | Cofilin_R |
| | ***Cloning of Tornado circRNA reporter plasmids*** | |
| PK352 | tcagaagaggatctggaagagtgagaattcacatttgcttctgacacaactgtgt | Tor_HBB_F |
| PK353 | cttgtcatcgtcgtccttgtagtccgtacgCATggtgtctgtttgaggttgctag | Tor_HBB_R |
| PK354 | tcagaagaggatctggaagagtgagaattcggtgtctgtttgaggttgctagtga | Tor_HBBinv_F |
| PK355 | cttgtcatcgtcgtccttgtagtccgtacgCATacatttgcttctgacacaactgtgt | Tor_HBBinv_R |
| PK356 | tcagaagaggatctggaagagtgagaattcTTGGGGGCGACACTCCAC | Tor_HCV_F |
| PK357 | cttgtcatcgtcgtccttgtagtccgtacgCATTGTTACGTTTGGTTTTTCTTTGAGGT | Tor_HCV_R |
| PK358 | tcagaagaggatctggaagagtgagaattcTGTTACGTTTGGTTTTTCTTTGAGGTTTAGGA | Torn_HCVinv_F |
| PK359 | cttgtcatcgtcgtccttgtagtccgtacgCATTTGGGGGCGACACTCC | Torn_HCVinv_R |
| PK360 | tcagaagaggatctggaagagtgagaattcTTAAAACAGCCTGTGGGTTGATCCC | Torn_CVB3_F |
| PK361 | cttgtcatcgtcgtccttgtagtccgtacgCATTGGTTTGCTGTATTCAACTTAACAATGAATTG | Torn_CVB3_R |
| PK362 | tcagaagaggatctggaagagtgagaattcGGTTTGCTGTATTCAACTTAACAATGAATTGTAATGT | Torn_CVB3inv_F |
| PK363 | cttgtcatcgtcgtccttgtagtccgtacgCATTTAAAACAGCCTGTGGGTTGATCC | Torn_CVB3inv_R |
| PK364 | tcagaagaggatctggaagagtgagaattcGTGTGGTGGGGGCGC | Torn_Chrdl1_F |
| PK365 | cttgtcatcgtcgtccttgtagtccgtacgCATCTTCTACTTTTTTCTCCTTCGAGCTACT | Torn_Chrdl1_R |
| PK366 | tcagaagaggatctggaagagtgagaattcCTTCTACTTTTTTCTCCTTCGAGCTACTGC | Torn_Chrdl1inv_F |
| PK367 | cttgtcatcgtcgtccttgtagtccgtacgCATGTGTGGTGGGGGCG | Torn_Chrdl1inv_R |
| PK368 | tcagaagaggatctggaagagtgagaattcCAGGCGTTGGGGGCG | Torn_Dlx1_F |
| PK369 | cttgtcatcgtcgtccttgtagtccgtacgCATCTCTTCTCGCGGGGT | Torn_Dlx1_R |
| PK370 | tcagaagaggatctggaagagtgagaattcCTCTTCTCGCGGGGTCTGG | Torn_Dlx1inv_F |
| PK371 | cttgtcatcgtcgtccttgtagtccgtacgCATCAGGCGTTGGGGGC | Torn_Dlx1inv_R |
| PK372 | tcagaagaggatctggaagagtgagaattcGCGCCCGCCCCT | Torn_Bcl2_F |
| PK373 | cttgtcatcgtcgtccttgtagtccgtacgCATCCTTCCCAGAGGAAAAGCAAC | Torn_Bcl2_R |
| PK374 | tcagaagaggatctggaagagtgagaattcCCTTCCCAGAGGAAAAGCAACG | Torn_Bcl2inv_F |
| PK375 | cttgtcatcgtcgtccttgtagtccgtacgCATGCGCCCGCCC | Torn_Bcl2inv_R |
| PK376 | tcagaagaggatctggaagagtgagaattcACTAGAACTCGCTGTAGTAATTCCAGC | Torn_Cmyc_F |
| PK377 | cttgtcatcgtcgtccttgtagtccgtacgCATTCGCGGGAGGCTGC | Torn_Cmyc_R |
| PK378 | tcagaagaggatctggaagagtgagaattcTCGCGGGAGGCTGCT | Torn_Cmycinv_F |
| PK379 | cttgtcatcgtcgtccttgtagtccgtacgCATACTAGAACTCGCTGTAGTAATTCCAGC | Torn_Cmycinv_R |
| PK380 | tcagaagaggatctggaagagtgagaattcGCCGGAAGGCCGCC | Torn_Cofilin_F |
| PK381 | cttgtcatcgtcgtccttgtagtccgtacgCATGTTTCCGGAAACGAAAGGGAG | Torn_Cofilin_R |
| PK382 | tcagaagaggatctggaagagtgagaattcGTTTCCGGAAACGAAAGGGAGACA | Torn_Cofilininv_F |
| PK383 | cttgtcatcgtcgtccttgtagtccgtacgCATGCCGGAAGGCCG | Torn_Cofilininv_R |
| PK384 | tcagaagaggatctggaagagtgagaattcttgatcttttaatcttcgttggccacaattaaaac | Torn_Hoxa9_F |
| PK385 | cttgtcatcgtcgtccttgtagtccgtacgCATtgcagtagcccgcg | Torn_Hoxa9_R |
| PK386 | tcagaagaggatctggaagagtgagaattctgcagtagcccgcgc | Torn_Hoxa9inv_F |
| PK387 | cttgtcatcgtcgtccttgtagtccgtacgCATttgatcttttaatcttcgttggccaca | Torn_Hoxa9inv_R |
| PK388 | tcagaagaggatctggaagagtgagaattcatcaggcaggatttacgactgg | Torn_A5_F |
| PK389 | cttgtcatcgtcgtccttgtagtccgtacgCATtgcttgatttgtggctcgc | Torn_A5_R |
| PK390 | tcagaagaggatctggaagagtgagaattctgcttgatttgtggctcgcg | Torn_A5inv_F |
| PK391 | cttgtcatcgtcgtccttgtagtccgtacgCATatcaggcaggatttacgactgg | Torn_A5inv_R |
